# Genomic structure predicts metabolite dynamics in microbial communities

**DOI:** 10.1101/2020.09.29.315713

**Authors:** Karna Gowda, Derek Ping, Madhav Mani, Seppe Kuehn

**Affiliations:** Department of Ecology and Evolution, University of Chicago, Chicago, IL 60637, USA; Center for the Physics of Evolving Systems, University of Chicago, Chicago, IL 60637, USA; Department of Physics, University of Illinois at Urbana-Champaign, Urbana, IL 61801, USA; Department of Engineering Sciences and Applied Mathematics, Northwestern University, Evanston, IL 60208, USA; Department of Molecular Biosciences, Northwestern University, Evanston, IL 60208, USA; NSF-Simons Center for Quantitative Biology, Northwestern University, Northwestern University, Evanston, IL 60208, USA

**Keywords:** microbial ecology, microbial communities, biological physics, statistical genomics, microbial community metabolism, denitrification, population dynamics, microbial interactions, community assembly, theoretical ecology

## Abstract

The metabolic function of microbial communities has played a defining role in the evolution and persistence of life on Earth, driving redox reactions that form the basis of global biogeochemical cycles. Community metabolism emerges from a hierarchy of processes including gene expression, ecological interactions, and environmental factors. In wild communities, gene content is correlated with environmental context, but predicting metabolic dynamics from genomic structure remains elusive. Here we show, for the process of denitrification, that community metabolism is predictable from the genes each member of the community possesses. Machine learning reveals a sparse and generalizable mapping from gene content to metabolite dynamics across a genomically-diverse library of bacteria. A consumer-resource model correctly predicts community metabolism from single-strain phenotypes. Our results demonstrate that the conserved impacts of metabolic genes can predict community function, enabling the prediction of metabolite dynamics from metagenomes, designing denitrifying communities, and discovering how genome evolution impacts metabolism.

## Introduction

The emergent metabolism of microbial communities plays an essential role in sustaining life on Earth, impacting global nutrient cycles (Falkowski et al., 2008; Canfield et al., 2010; Stein and Klotz, 2016), wastewater treatment (Lu et al., 2014) and human health (Subramanian et al., 2014). An elusive challenge in microbial ecology is understanding how emergent community metabolism is determined by the taxonomic and genomic structure of a community (Widder et al., 2016; Louca, Polz, et al., 2018). Addressing this challenge requires mapping the genotypes of each community member to metabolic phenotypes, and then deciphering how complex interactions between distinct populations, which depend on extracellular metabolites (Lilja and Johnson, 2016), abiotic factors (Ward et al., 2006), cooperation (Cordero et al., 2012), and higher-order effects (Sanchez-Gorostiaga et al., 2019; Mickalide and Kuehn, 2019), contribute to the collective. Solving this structure-function problem is critical for functionally interpreting community gene content (Anantharaman et al., 2016), designing synthetic communities (Shou et al., 2007), and elucidating the evolutionary principles of community metabolism (Molina and Nimwegen, 2009; Sela et al., 2019).

Recent studies hint that gene content may be predictive of emergent metabolic function at the community level. Sequencing studies of environmental and host-associated communities show that, while abundances of individual taxa can be highly variable (Louca, Polz, et al., 2018), the genes or pathways a community possesses are stable across communities in similar environments (Louca, Polz, et al., 2018; Huttenhower et al., 2012). This suggests that gene content is a conserved feature of natural communities (Burke et al., 2011). Additional studies have extended this insight by demonstrating that the functional gene content of a community correlates with local metabolite concentrations (Jones and Hallin, 2010; Fierer et al., 2012; Louca, Parfrey, et al., 2016). However, a major limitation of sequencing studies of natural communities is that we cannot easily disentangle genomic and environmental impacts on community structure and function. For example, the observed correlation between local metabolite concentrations and gene content in the global ocean microbiome (Louca, Parfrey, et al., 2016), may arise because community gene content determines metabolite fluxes, or alternatively because exogenously controlled metabolite fluxes determine community metagenomic structure.

Therefore, the key to unambiguously mapping gene content to metabolite dynamics is laboratory experiments with genomically diverse communities where the environmental context can be controlled. To attempt this, several studies have taken an enrichment culture approach where communities from the wild are grown in simple nutrient conditions in the laboratory (Datta et al., 2016; Goldford et al., 2018). These experiments have revealed conserved metabolic traits in members of the assembled communities, such as the ability to degrade chitin in the primary colonizers of chitin particles (Datta et al., 2016), or family-level conservation of traits governing glucose utilization and subsequent cross-feeding (Goldford et al., 2018). However, these studies have not successfully mapped the gene content of assembled communities to their metabolic function, in large part because the enrichment process in simple nutrient conditions dramatically reduces the diversity of the wild community (Jiao et al., 2016). A loss of diversity means that much of the variation in functional gene content is lost during enrichment, and with it variation in community function (Jiao et al., 2016; Conthe et al., 2018; Goldford et al., 2018). So while enrichment experiments permit precise control over the environment and reveal conserved features of assembled communities, they reduce variation in genomic structure and metabolic function of the assembled communities, making it challenging to determine how variation in structure impacts function.

Here we address the challenge of mapping gene content to metabolite dynamics by quantifying the flux of metabolites in an ensemble of genomically-diverse communities composed of non-model organisms. We used bacterial denitrification, an essential metabolic process in the global nitrogen cycle, as a model metabolic function that is performed by diverse and culturable bacterial taxa (Lycus et al., 2017). We isolated an ensemble of denitrifiers and measured the dynamics of metabolite consumption and production for each isolate under controlled conditions. We then parameterized metabolite dynamics using a simple consumer-resource model. The genomic diversity of the ensemble of isolates enabled a statistical learning approach to mapping gene content to consumer-resource model parameters, which resulted in a sparse and generalizable mapping of gene presence and absence to metabolic phenotypes. Finally, the consumer-resource model captured interactions between strains mediated by resource competition, yielding predictions for community-level metabolite dynamics, which we verified experimentally.

## Results

### Denitrification as a model metabolic process

We used denitrification as a model metabolic process (Fig. 1A) because it is performed by diverse bacterial taxa, it is well-characterized at the molecular level, it is a collective process, and the relevant metabolites are readily quantifiable (Zumft, 1997). Because denitrifiers are easily isolated and cultured (Lycus et al., 2017), we can capture substantial genomic diversity in an ensemble of natural isolates.

**Figure 1:**
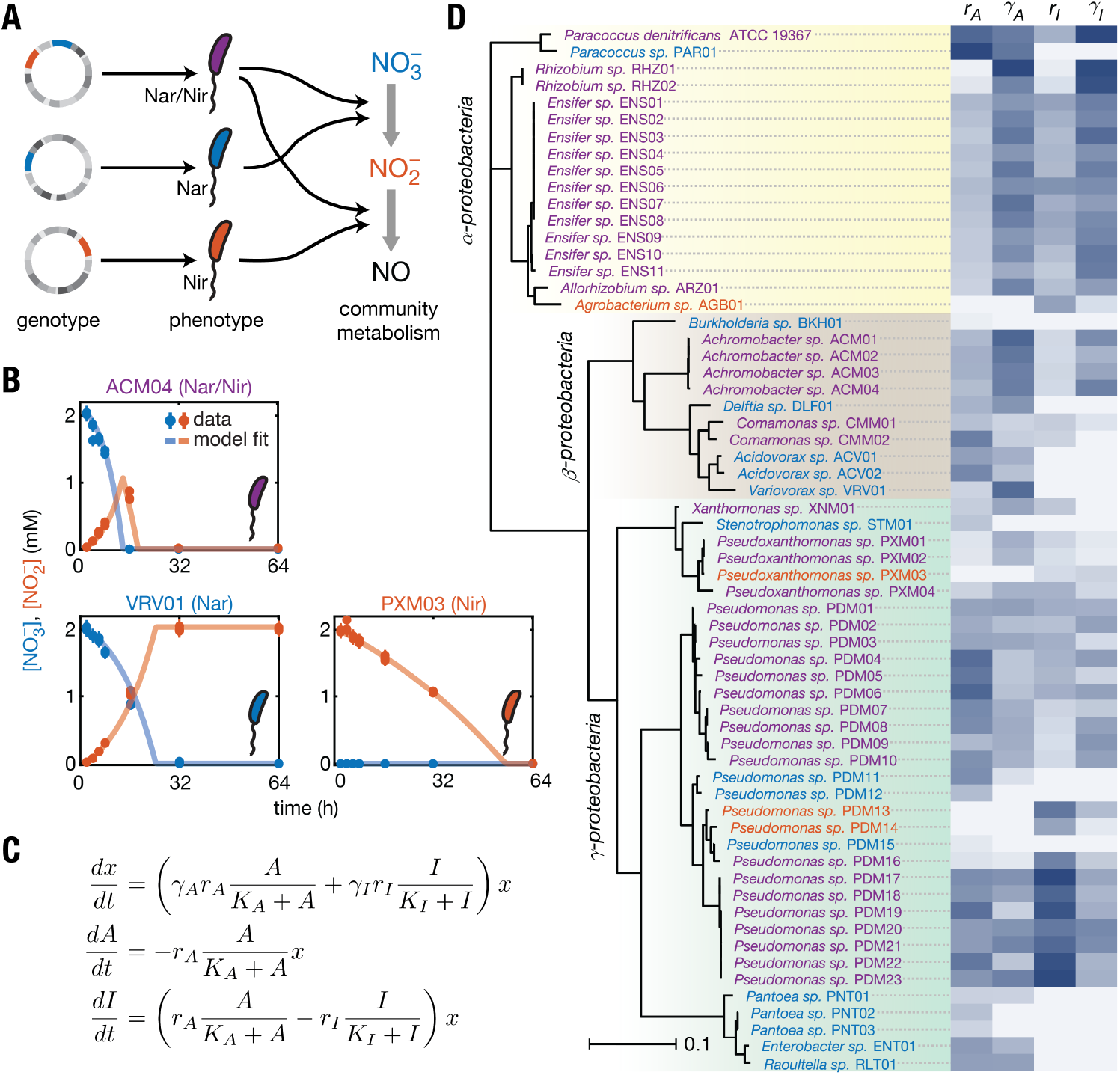
Quantifying nitrate and nitrite dynamics in an ensemble of denitrifiers to map genomic structure to metabolic function. (**A**) A roadmap for relating genomic structure to community metabolic function across a diverse library of bacteria. First, mapping the genotypes (circles) of individual strains to their metabolic phenotypes (straight arrows) and then combining the metabolic activity of each strain in a community to predict collective metabolite dynamics (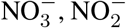, right). (**B**) Example batch culture metabolite dynamics for Nar/Nir (purple), Nar (blue), and Nir (red) isolates. Nitrate (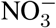, blue points) and nitrite (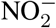, red points) dynamics are measured at logarithmically-spaced intervals (circles) via sampling and colorimetric assay (see Methods). Abundances are only measured at the final time point. Curves show fits to a consumer-resource model shown in panel C. (**C**). A consumer-resource model of nitrate and nitrite reduction by each strain describes abundances (*x*), nitrate concentration (*A*), and nitrite concentration (*I*) in time. The model is parameterized by reduction rates *r_A_* and *r_I_* and yields *γ_A_* and *γ_I_*, for growth on nitrate and nitrite respectively. The affinity parameters (*K*_*_) were not well-constrained by the data and were fixed for all strains in the library (see Supplemental Information). (**D**) Phylogenetic tree and normalized consumer-resource parameters for 62 denitrifying strains (61 isolates and the model denitrifier *Paracoccus denitrificans*). Phylogenetic tree constructed using the 16S rRNA gene, and scale bar indicates the estimated number of substitutions per site. Darker colors indicate larger values of the normalized parameters. Each isolate was assigned a unique identifier. Phenotypic parameters measured across diverse isolates constitutes a dataset for relating genomic diversity to metabolite dynamics. See also Figs. S1–S7.

Denitrification is a form of anaerobic respiration whereby microbes use oxidized nitrogen compounds as electron acceptors, driving a cascade of four successive reduction reactions, 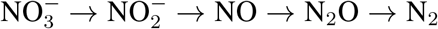 (Zumft, 1997). As a biogeochemical process, denitrification is essential to nitrogen cycling at a global scale through activity in soils, freshwater systems, and marine environments (Seitzinger et al., 2006), and impacts human health through activity in wastewater treatment plants (Lu et al., 2014) and in the human gut (Irrazabal et al., 2014). The process is performed by taxonomically-diverse bacteria (Graf et al., 2014) that are typically facultative anaerobes. The denitrification pathway is known to be modular, with some strains performing all four steps in the cascade, and others performing one or a nearly arbitrary subset of reduction reactions (Lycus et al., 2017). Denitrification in nature is therefore a collective process, where a given strain can produce electron acceptors that can be utilized by other strains (Lilja and Johnson, 2016).

We focused experimentally on the first two steps of denitrification: the conversion of nitrate 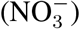 to nitrite 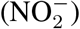 and subsequently nitric oxide (NO) (Fig. 1A). Nitrate and nitrite are soluble, enabling high throughput measurements of metabolite dynamics. In order to obtain a genomically-diverse ensemble of non-model organisms, we isolated 61 bacterial strains spanning *α*-, *β*-, and *γ*-proteobacteria from local soils using established techniques (Methods). Each strain was obtained in axenic culture and was characterized as performing one or both of the first two steps of denitrification in a chemically-defined, electron acceptor-limited medium containing a single non-fermentable carbon source (succinate). Each of these strains was therefore classified into one of three possible phenotypes (Fig. 1A): (1) Nar/Nir strains that perform both nitrate and nitrite reduction 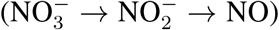, (2) Nar strains that perform only nitrate reduction 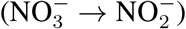, and (3) Nir strains that perform only nitrite reduction 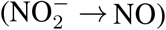. In addition to these 61 isolates, our strain library also included the model Nar/Nir strain *Paracoccus denitrificans* (ATCC 19367).

### Parameterizing metabolite dynamics

We first set out to quantify the metabolic phenotypes of each isolate in our diverse strain library. We focused our efforts on quantifying the dynamics of the relevant metabolites, nitrate and nitrite. To accomplish this, strains were inoculated at low starting densities into 96-well plates containing chemically-defined medium with either nitrate or nitrite provided as the sole electron acceptor, and then incubated under anaerobic conditions. Small samples (10 μL) were then taken at logarithmically-spaced time intervals over a period of 64 h and assayed for nitrate and nitrite concentrations (Methods, Fig. S1 and S2). At the end of the time course, optical density was assayed. The measurement resulted in a time series of nitrate and nitrite production/consumption dynamics in batch culture (points, Fig. 1B).

To parameterize the metabolite dynamics of each strain within a common framework, we utilized a consumer-resource model, which explicitly relates the growth of each strain to the dynamics of metabolite production and consumption (Fig. 1C). For each strain in monoculture, we parameterized the consumer-resource model using measured denitrification dynamics across a range of initial cell densities and nitrate/nitrite concentrations (Supplemental Information, Fig. S3 and S4). The model allowed us to quantitatively describe the phenotype of each strain in the library using at most four parameters: *r_A_* and *r_I_*, which capture rates of nitrate and nitrite reduction, and and *γ_A_*, and *γ_I_* which describe yields for nitrate and nitrite, respectively. Substrate affinities (*K*_*_) were fixed to a small value since these parameters were not well constrained by the data (Supplemental Information, Fig. S5). The models for Nar and Nir strains correspond to setting *γ_I_* = 0 or *γ_A_* = 0, respectively. Yields (*γ*_*_) were inferred using endpoint optical density measurements, and rates (*r*_*_) were inferred by fitting the observed nitrate and nitrite dynamics to the consumer-resource model (Fig. 1C). Remarkably, with the exception of a small number of strains that were excluded from the library (Supplemental Information, Fig. S6), a single set of parameters quantitatively described metabolite dynamics for each strain across a range of initial cell densities and nitrate/nitrite concentrations (Supplemental Information, Fig. S4).

Fitting our consumer-resource model to data for each strain yielded a quantitative description of the dynamic metabolic phenotype of each strain in the library (Fig. 1B and D). We observed large variability between taxa, with coefficients of variation for both rates (*r_A_, r_I_*) and yields (*γ_A_, γ_I_*) around 60%. We also observed some patterns of phylogenetic conservation, for example *α*-proteobacteria produced generally higher yields than *β*-or *γ*-proteobacteria, and a clade of *Pseudomonas* sp. isolates showed consistently higher rates of nitrite reduction than most other strains (Fig. 1D). Despite these patterns, the prevalence of each of the three phenotypes is not strongly dependent on phylogeny, with each phenotype present across the tree (Fig. 1D). The latter observation is consistent with pervasive horizontal gene transfer of denitrifying enzymes (Heylen et al., 2006; Jones, Stres, et al., 2008). Finally, we did not observe a trade-off between rates and yields (Fig. S7).

### Predicting metabolite dynamics from genomes

Mapping genomic structure to metabolic function in communities requires understanding the phenotypic impacts of genomic variation at the level of individual genomes. The problem translates into understanding how genomic variation across the strains in our library gives rise to variation in the quantitative metabolic traits (Fig. 1D). One common approach to the problem of relating genomes to metabolite dynamics is constraint-based modeling. Constraint based models infer or measure the set of all metabolic reactions performed by a given organism and predict growth and metabolite fluxes assuming the metabolic network is in steady state and subjected to biologically motivated constraints (Orth et al., 2010). Constraint-based methods have found some success in predicting collective metabolism from genomes (Klitgord and Segrè, 2011; Mori et al., 2016; Harcombe et al., 2014), but these methods require significant manual refinement (Norsigian et al., 2020), complicating the prospect of making predictions from the genomes of non-model organisms. As a result, successfully constructing constraint-based models of denitrification for all strains in our library is a daunting task.

We took an alternative approach to the problem of mapping genomes to metabolite dynamics. We asked whether whether the variation in metabolic phenotypes across strains in our library can be quantitatively predicted simply from knowledge of the genes possessed by each strain. Our conjecture was motivated by the observation that gene content at the community-level correlates strongly with environmental variation in natural communities (Louca, Parfrey, et al., 2016), and that the statistics of gene presence and absence capture significant functional information (Schober et al., 2019). Therefore, we took a simple statistical learning approach to predicting the rates and yields measured across our library of isolates (Fig. 1D) from genomic structure. Specifically, we predicted the consumer-resource model parameters from gene presence and absence alone. We performed whole genome sequencing on all 62 strains in the library. Then we assembled and annotated each genome (Methods), and determined the complement of 17 denitrification-related genes possessed by each strain, exploiting the fact that the molecular and genetic basis of denitrification is well-understood (Zumft, 1997). We identified not only the reductases that perform the reduction of the oxidized nitrogen compounds, but also the sensors/regulators (Rodionov et al., 2005) and transporters (Moir and Wood, 2001) known to be involved in denitrification (Methods). The presence and absence of each gene (or set of genes encoding proteins that form a complex) in each genome is presented in Fig. 2A. Patterns of gene presence and absence agree well with known features of the denitrification pathway, including the mutual exclusion of the two reductases performing nitrite reduction (*NirS* and *NirK*) (Jones, Stres, et al., 2008; Jones and Hallin, 2010). Further, in almost all cases strains possessing nitrate and/or nitrite reductase performed the associated reactions in culture (with the only exception being the Nar strain *Acidovorax* sp. ACV01, which possesses both nitrate and nitrite reductase). This is in agreement with previous work demonstrating that bacterial genomes lose nonfunctional genes due to streamlining (Lynch, 2006).

**Figure 2:**
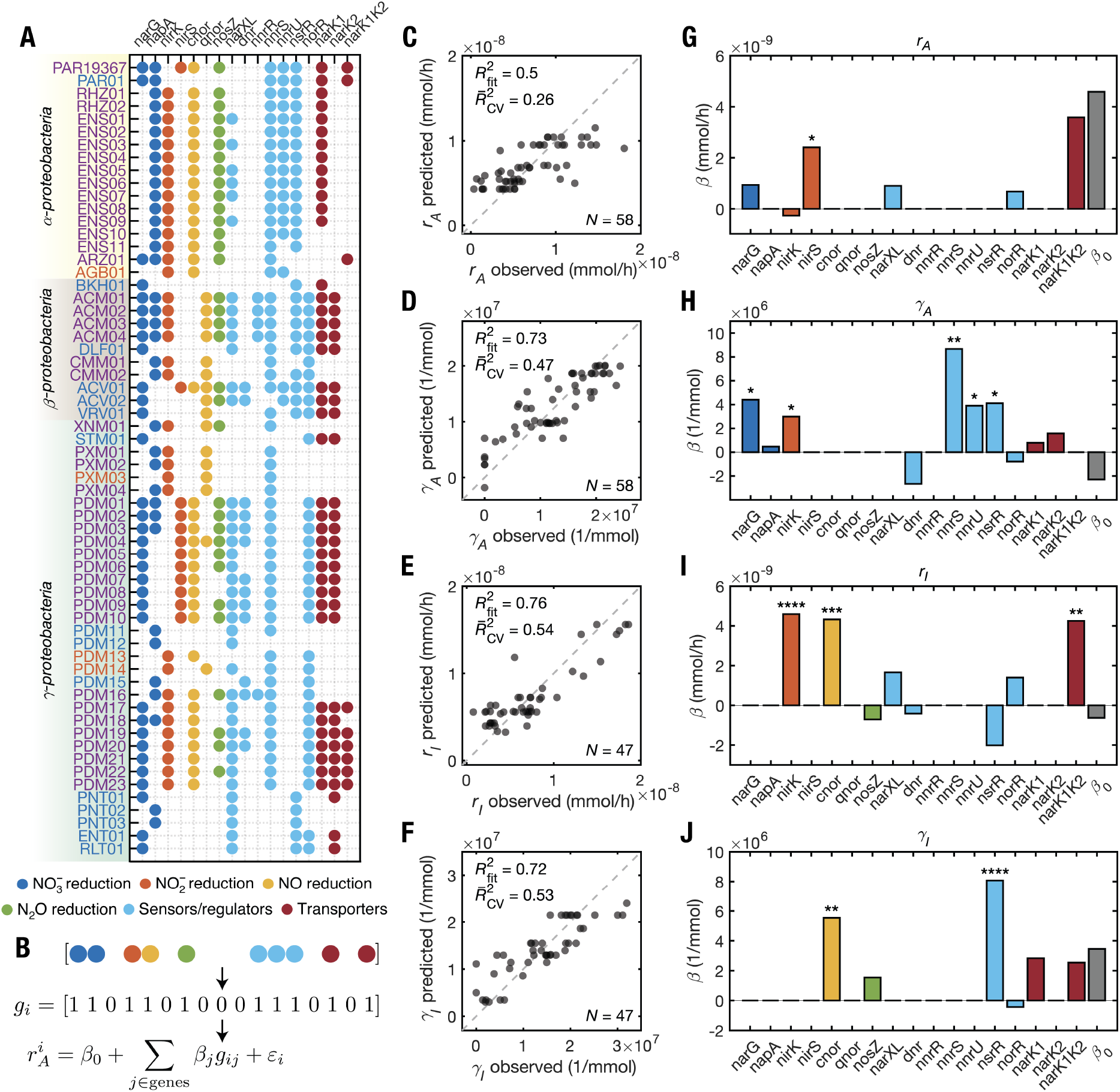
A statistical mapping from gene presence and absence to metabolite dynamics of individual strains. (**A**) The presence and absence of genes in the denitrification pathway for the 62 denitrifying strains in our library. Strain identifiers correspond to those given in Fig. 1D. The color of each circle corresponds to the known gene function as indicated in the legend below. (**B**) Observed consumer-resource phenotypic parameters for each strain (e.g., nitrate reduction rate *r_A_*, Fig. 1D) were linearly regressed against gene presence and absence via *L*_1_-regularized regression, resulting in regression coefficients *β_j_* for each gene *j*, an intercept *β*_0_, and a noise term *ϵ_i_* for each observation *i*. Coefficient *β_j_* captures the impact of possessing gene *j* on the corresponding phenotypic parameter. Independent regressions were performed for each phenotypic parameter. **C, D, E, F** show predicted values of *r_A_, γ_A_, r_I_*, and *γ_I_* respectively plotted against measured values. The dashed line indicates perfect agreement between observations and predictions. The in-sample coefficients of determination for these data 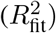 and the out-of-sample coefficients of determination estimated via iterated cross-validation 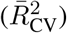 are shown. *N* indicates the number strains in each regression. Strains which do not perform a particular reaction were omitted from the corresponding regression (e.g., Nir strains were excluded from the regression for *r_A_*). **G, H, I, J** show estimates of *β_j_* and *β*_0_ for *r_A_, γ_A_, r_I_*, and *γ_I_* respectively. Asterisks indicate significance level for each *β* (*: *p* ≤ 0.05, **: *p* ≤ 10^-2^, ***: *p* ≤ 10^-3^, ****: *p* ≤ 10^-4^; see Supplemental Information). See also Figs. S8-S14.

Next we showed that the presence and absence of denitrification genes in each strain was sufficient to quantitatively predict metabolite dynamics in monoculture. Specifically, we constructed a linear regression where the measured phenotypic parameters of our consumer-resource model were predicted on the basis of gene presence and absence (Fig. 2B). The regression coefficients for each gene in the pathway quantify the impact of the presence of the gene on a given phenotypic parameter. We used *L*_1_-regularized regression (least absolute shrinkage and selection operator, LASSO) to avoid overfitting (Supplemental Information, Fig. S8 to S10), performing independent regressions for each of the phenotypic parameters in our consumer-resource model. LASSO yielded sparse regression models, revealing that presence and absence of a small set of genes is highly predictive of the phenotypic parameters for all strains in our library (Fig. 2C to J). The in-sample coefficients of determination 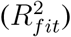 of our regressions were between 0.50 and 0.76 depending on the phenotypic parameter. Crucially, our regression approach generalized out-of-sample, as determined by iterated cross-validation (Supplemental Information, Fig. S9), albeit with slightly lower predictive power (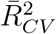 between 0.26 to 0.54). Since, in general, traits may exhibit phylogenetic correlation (Martiny et al., 2015), and our library contains a few clades comprising very closely related strains (e.g., ENS01–08, PDM20– 23, Fig. 1D), we considered whether our regression utilized phylogenetic correlations in gene presence and absence and denitrification phenotypic parameters to achieve predictive power. We investigated this by collapsing clades containing strains with identical 16S rRNA sequences down to a single randomly-selected representative, and performing regressions again on this reduced set of strains. For these regressions we found that the predictive power and coefficients were similar to those for the full dataset (Supplemental Information, Fig. S11), supporting the claim that our regression is not simply detecting phylogenetic correlations between traits and genotypes. Altogether these results demonstrate that, across a diverse set of natural isolates, knowledge of the genes a denitrifying strain possesses is sufficient to accurately predict the rates and biomass yields of that strain on nitrate and nitrite.

Our regression approach leveraged biological knowledge of the denitrification pathway to predict metabolite dynamics, in effect presuming that denitrification gene content is the best genomic feature for prediction. To investigate whether this assumption is correct, we asked whether other genomic properties could better predict metabolite dynamics. First, we tested the predictive capability of sets of randomly selected genes. We chose sets of 17 random genes that were not strongly correlated with any denitrification genes, but retained the same marginal frequency distribution as denitrification genes in the population. We found that regressions using these randomly-selected genes have, on average, much less predictive power than regressions using the denitrification genes (Supplemental Information, Fig. S12). We note that this result provided further evidence that regressions on denitrification gene presence and absence are not simply detecting phylogenetic correlations, since random genes would be expected to perform equally well on average if phenotypes were simply determined by phylogeny. Second, we tested whether 16S rRNA copy number or genome size improves the predictive ability of denitrification gene presence and absence regressions. 16S rRNA copy number has been observed to correlate positively with maximal growth rate in nutrient rich conditions (Roller et al., 2016; Li et al., 2019), and smaller genomes are associated with faster growth (Lynch, 2006; Li et al., 2019). We found that including these predictors in our regressions does not meaningfully improve their predictive ability or alter the inferred coefficients (Supplemental Information, Fig. S13 and S14). In summary, our statistical analyses provided evidence that denitrification gene presence and absence outperforms arbitrary sets of genes and coarse genomic features.

Why does gene presence and absence alone hold such strong predictive power for metabolite dynamics, and why did the regression select specific genes in the denitrification pathway as informative predictors? We propose that by quantifying metabolic phenotypes in terms of rates and yields, we captured the salient features of the metabolic process for each strain, which allowed the regression to succeed by exploiting the conserved impacts of specific genes on these metabolic phenotypes. To investigate this claim we examined the regression coefficients in the context of what is known about the denitrification pathway. We found that in many cases the sign and magnitude of the regression coefficients agree qualitatively with known properties of the associated enzymes. Previous comparisons between membrane-bound and periplasmic nitrate reductases (encoded by *narG* and *napA*, respectively) in multiple bacterial species showed that the membrane-bound enzyme exhibits higher nitrate reduction activity *in vitro* than the periplasmic enzyme (Table S1). This accords with the large positive coefficient for *narG* we observed in the nitrate reduction rate regression (Fig. 2G). Similarly, in the nitrite reduction rate regression we observed a large positive coefficient for the gene encoding the copper-based nitrite reductase (*nirK*) (Fig. 2I), which in previous studies showed markedly higher activity *in vitro* (Table S2) and *in vivo* (Gloekner et al., 1993) compared to the alternate nitrite reductase enzyme encoded by *nirS*. Further, our regression coefficients showed larger contributions of *narG* versus *napA* to yield on nitrate (Fig. 2H), and similarly *cnor* versus *qnor* to yield on nitrite (Fig. 2J). Both these observations are consistent with the fact that the genes encoded by *narG* and *cnor* contribute more to the proton motive force (and therefore to ATP generation) than their alternatives, *napA* and *qnor*, respectively (Ferguson and Richardson, 2004). Finally, the transporter encoded by the gene *narK1K2* is a fusion of the nitrate/H^+^ symporter *NarK1* and the nitrate/nitrite antiporter *NarK2*, the latter of which is crucial for exchanging nitrate and nitrite between the cytoplasm and periplasm during denitrification when the membrane-bound nitrate reductase is utilized. In *Paracoccus denitrificans*, this fusion has been shown to have substantially higher affinity for nitrate than *NarK2* alone, resulting in higher growth rates under denitrifying conditions (Goddard et al., 2008). Remarkably this agrees with what we found in the nitrate and nitrite reduction rate regressions, where we observed large positive contributions of *narK1K2* (Fig. 2G and I). Taken together, these observations suggest that the regressions exploited mechanistic aspects of the denitrification process to predict metabolite dynamics. However, for many coefficients in our regression, notably regulators, there is no clear interpretation, and definitive proof that these coefficients are mechanistically informative will require genetic manipulation of diverse bacteria.

Our statistical approach took two important steps towards mapping genomic structure to metabolic dynamics at the single-strain level. First, by making quantitative measurements in the laboratory, we removed the confounding environmental factors present in sequencing and metabolomic studies of natural communities to reveal that gene content has a conserved impact on dynamical metabolic phenotypes. Second, our results suggest that a statistical approach could be used to discover the key genomic features of pathways that determine other metabolic phenotypes, complementing direct genetic interrogation of model organisms (Nichols et al., 2011). Further, our predictions of metabolic phenotypes from genomes apply across a range of initial conditions and generalized well out-of-sample, suggesting that this approach can predict metabolite dynamics in a variety of settings for strains where only genome sequence data are available. These insights were made possible by parameterizing metabolic phenotypes across a genomically-diverse strain library of non-model organisms, thereby exploiting genomic variation to learn the mapping from genotype to metabolic phenotypes.

### Predicting metabolite dynamics in communities

Predicting community metabolite dynamics from genomic structure next requires mapping single-strain phenotypes to collective behavior. The consumer-resource modeling formalism (Fig. 1C) we used to parameterize metabolite dynamics for each strain allows us to make quantitative predictions for metabolite dynamics in communities of multiple taxa. Since phenotypic parameters were sparsely-encoded by the genomes of each strain (Fig. 2), predicting community metabolite dynamics from the consumer-resource model would provide a direct mapping from gene content to community metabolism. Therefore, we extended to our modeling formalism to *N*-strain communities by adding the rate contributions of each strain to the dynamics of nitrate and nitrite (Fig. 3B, Supplemental Information). This model assumes that strains interact only via cross-feeding and resource-competition for electron acceptors. This “additive” model also assumes that the rates and yields on nitrate and nitrite for strains in pair-culture are the same as in monoculture. As a result, the model provides predictions for *N*-strain community metabolite dynamics without any free parameters.

**Figure 3:**
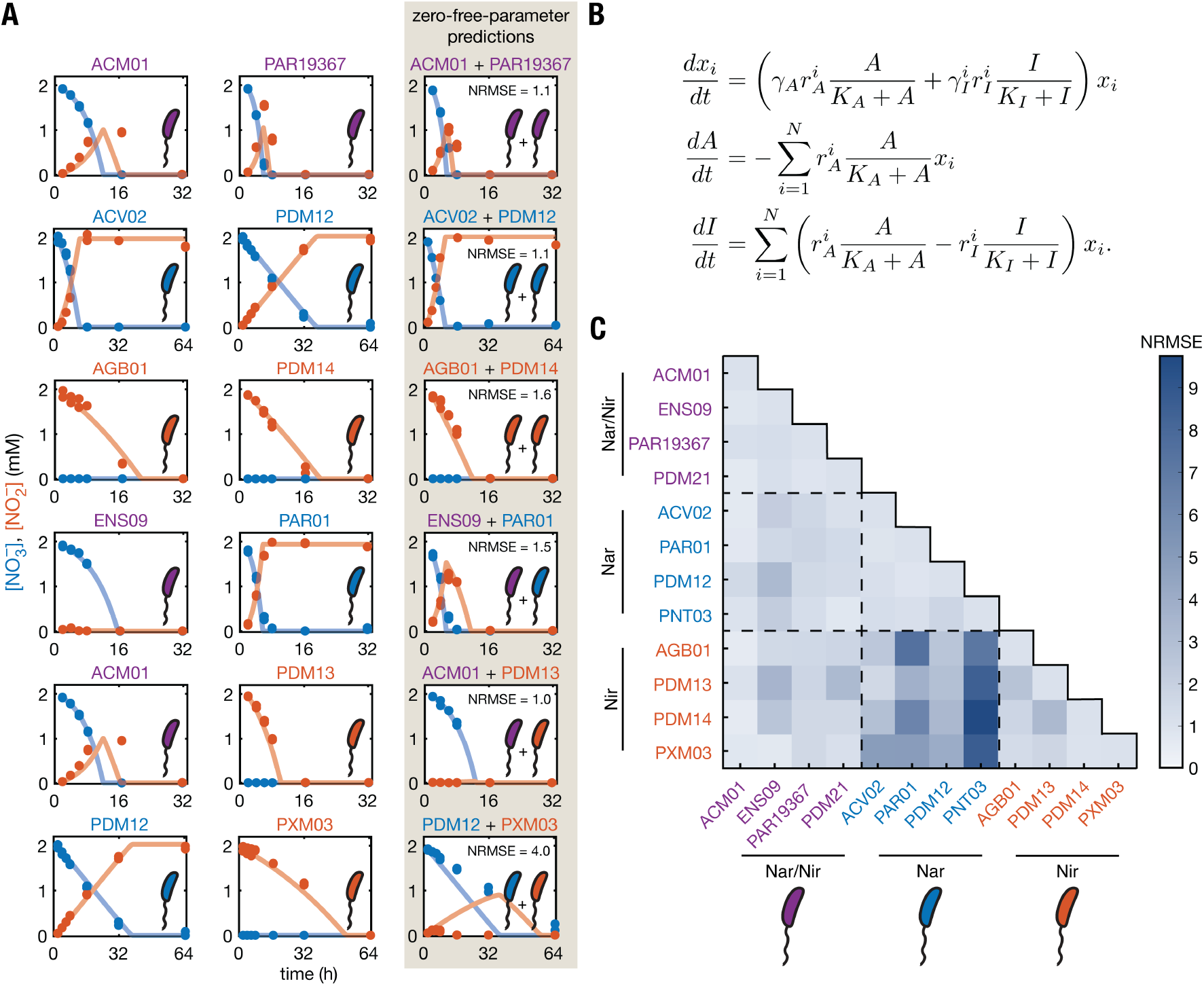
Metabolite dynamics in two-strain communities are predictable from monocultures. (**A**) Examples of pair-culture dynamics for all combinations of the three denitrification phenotypes (Nar/Nir, purple; Nar, blue; Nir; red). The first two columns show metabolite dynamics for each of two strains cultured individually. The third column shows the metabolite dynamics for pair-cultures of the two strains (points) with zero free parameter predictions using the consumer-resource model (curves, see panel **B** for model). Errors in pair-culture predictions are shown in each panel in the third column as quantified by the normalized root-mean-square error (NRMSE). We define 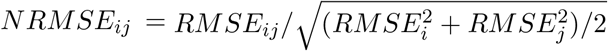, where RMSE*_ij_* is the root-mean-square error between model predictions and observed metabolite concentrations of strains *i* and *j* in pair-culture, and RMSE_*i*_ and RMSE_*j*_ are the RMSEs of strains *i* and *j* in monoculture. NRMSE values between 0 and 2 indicate predictions of pair-culture metabolite dynamics of similar quality to the corresponding monocultures. (**B**) An *N*-strain consumer-resource model (based on Fig. 1C) was used to predict pair-culture metabolite dynamics (*N* = 2). *A* and *I* are nitrate and nitrite concentrations respectively. *x_i_* denotes abundance of strain *i* with parameters 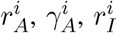 and 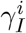, which were determined from monoculture experiments (Fig. 1D). The *K*_*_ values are fixed for all strains. (**C**) A matrix of NRMSE values quantifying the quality of model predictions for all pairs of 12 strains: 4 Nar/Nir, 4 Nar, 4 Nir as indicated left and below. Strain names correspond to Figs. 1D and 2A. Only Nar + Nir communities are poorly predicted by the consumer-resource model (permutation test, *p* < 10^-5^, Fig. S16, Figure 4). See also Figs. S15 and S16.

We first tested the ability of this approach to predict metabolite dynamics in all pair combinations of 12 strains from our library (4 Nar/Nir, 4 Nar, 4 Nir). We assembled communities in 96-well plates containing chemically-defined medium and sampled over a 64 h period to measure concentrations of nitrate and nitrite (Methods). Remarkably, we found that the additive model accurately predicted the metabolic dynamics for most 2-strain communities (Fig. 3, Fig. S15 and S16). Specifically, the third column of Fig. 3A shows the zero-free-parameter predictions (lines) of denitrification dynamics in 2-strain communities, which agreed well with measurements (points). The 2-strain community predictions include non-trivial dynamics such as two Nar strains exhibiting faster nitrate reduction as a collective or a transient increase nitrite in a Nar/Nir + Nar community.

We quantified the quality of the additive model predictions by computing a normalized root-mean-square error (NRMSE, see caption of Fig. 3). NRMSE in the range 0–2 indicates predictions in 2-strain communities that are similar in quality to the fits of the constituent monocultures. We found that most 2-strain communities have low NRMSE, indicating that our model successfully predicted metabolite dynamics in most cases, given only knowledge of the monoculture rates and yields for each strain. The success or failure of the model depended on the phenotypes of the strains present. The model successfully predicted 2-strain metabolite dynamics for most types of communities (e.g., Nar/Nir + Nar or Nar + Nar) but failed only in the case where Nar strains were cultured with Nir strains (Fig. 3A and C, Fig. S17). We speculate that the failure of the model to predict metabolite dynamics in Nar + Nir communities was caused by excretion of nitric oxide by the Nir strain, which can be cytotoxic to strains that do not express nitric oxide reductase (Braun and Zumft, 1991), and may consequently slow Nar strain growth. Although both Nar/Nir and Nir strains are capable of generating extracellular nitric oxide, Nir isolates have been observed in a previous study to transiently generate nitric oxide at higher concentrations (Lycus et al., 2017), possibly explaining why the 2-strain additive model fails only to predict Nar + Nir communities.

We next asked whether information from monocultures also successfully predicted metabolite dynamics in 3-strain communities. We applied the additive model to predicting the nitrate and nitrite dynamics in 81 random combinations of 3 strains from the 12-strain subset. In communities that did not contain a Nar + Nir pair (e.g., Fig. 4A), we found that prediction accuracy was high (grey points, Fig. 4B, Fig. S18). This again indicated that in most combinations of phenotypes, community dynamics were predictable from consumer-resource parameters for each strain in the community. However, in communities that contained a Nar + Nir pair, predictions were relatively poor (yellow points, Fig. 4B, Fig. S18), suggesting that interactions between Nar and Nir phenotypes that were not captured in the additive model were again driving low prediction accuracy.

**Figure 4:**
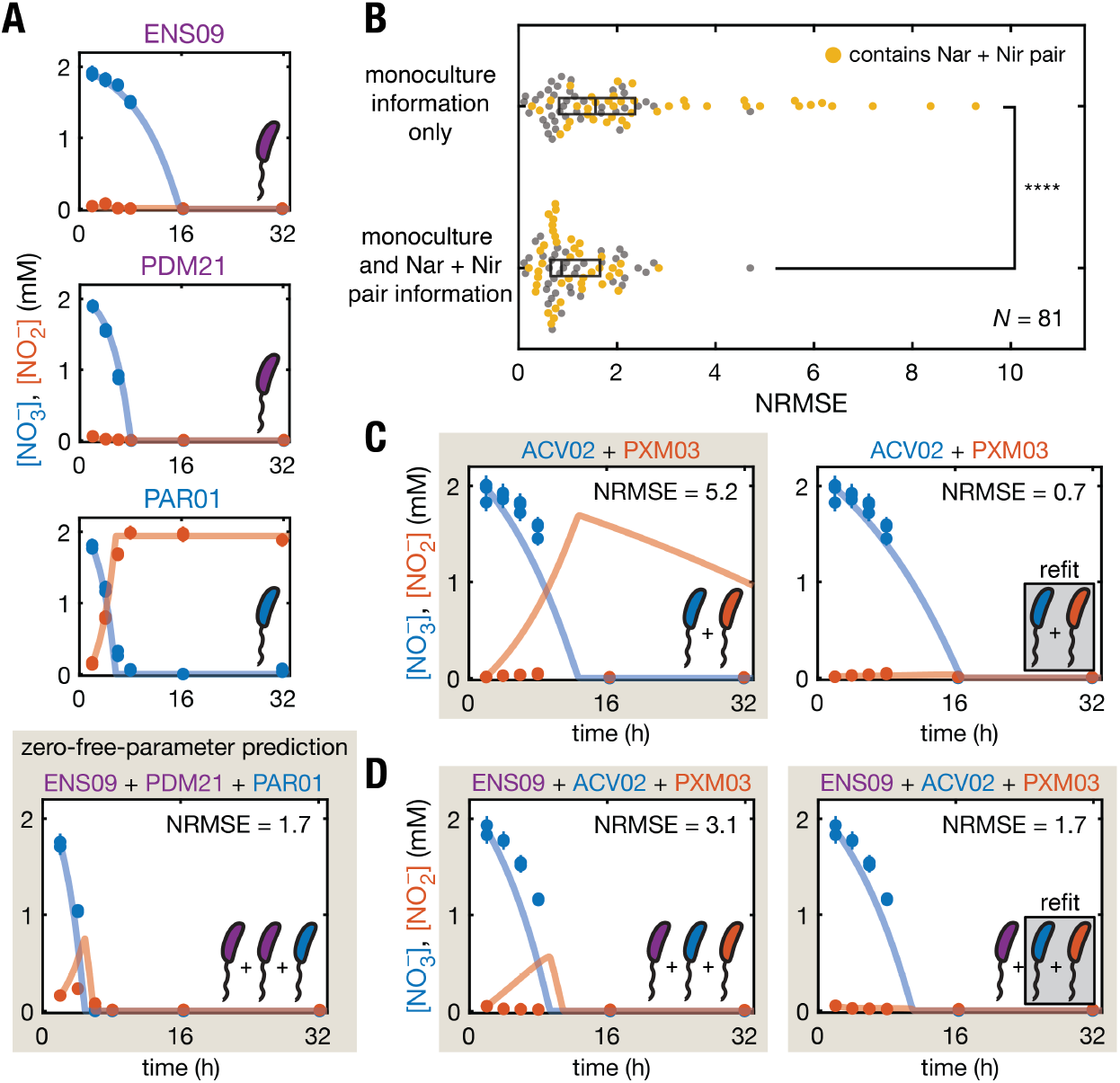
Metabolite dynamics in 3-strain communities are predictable. The consumer-resource model provides predictions for metabolite dynamics in communities of three strains, and these predictions were verified experimentally. (**A**) Metabolite dynamics for an example 3-strain (Nar/Nir + Nar/Nir + Nar) community. The first three panels show metabolite dynamics for each strain cultured individually, and the fourth panel shows the metabolite dynamics of the 3-strain community. Curves show the prediction of the consumer-resource model (Fig. 3B). (**B**) NRMSE (see Fig. 3 caption) values quantifying quality of consumer-resource model predictions for 3-strain communities. Orange and (gray) points denote 3-strain communities that do (do not) contain a Nar + Nir pair. Nar + Nir pair-culture dynamics are poorly predicted by the model (Fig. 3C) and result in high NRMSE in 3-strain communities containing Nar + Nir pairs (compare orange and gray points). Top and bottom scatter plots compare predictions from a consumer-resource model using only monoculture data to a coarse-graining approach that describes Nar + Nir pairs as modules within the 3-strain community (described in **C-D**). The coarse-graining approach improves the 3-strain community predictions (*t*-test, **** denotes *p* = 7 × 10^-6^). (**C**) Metabolite dynamics for an example Nar + Nir pair, where curves in the left panel show the prediction of the consumer-resource model using only parameters fit to monocultures, and curves in the right panel show the results of refitting the reduction rates (*r_A_, r_I_*) to Nar + Nir pair-culture data but leaving yields (*γ_A_, γ_I_*) fixed to monoculture values. (**D**) Metabolite dynamics for a 3-strain community containing a Nar/Nir strain and the Nar + Nir pair shown in panel **C**. Curves in the left panel show the prediction of the consumer-resource model using parameters inferred from monoculture experiments for each strain, and curves in the right panel show the prediction when the Nar + Nir pair is treated as a module with rate parameters refit from pair-culture data (right panel in C). Note the reduction in NRMSE due to the coarse-graining of the Nar + Nir pair. Panels in beige denote zero free parameter predictions. See also Figs. S17-S19.

To address the impact of interactions between Nar and Nir strains not accounted for by our additive model in 3-strain communities, we took a coarse-graining approach. We asked whether the collective metabolism of Nar + Nir pairs could be treated as modules within 3-strain communities. To accomplish this we re-fitted nitrate and nitrite reduction rates (*r_A_, r_I_*) to pairculture data for each Nar + Nir pair, leaving yields fixed (Fig. 4C, Supplemental Information, Fig. S19). This resulted in effective nitrate and nitrite reduction rates 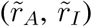 for each Nar + Nir pair. We then used these rates to make predictions for 3-strain communities that included a Nar + Nir pair (e.g., Fig. 4D). For 3-strain communities that included multiple Nar + Nir pairs (e.g., Nar + Nar + Nir), we developed simple rules for determining the effective rates from the rates for each Nar + Nir pair present (Supplemental Information). We found that the metabolite dynamics in 3-strain communities containing Nar + Nir pairs were quantitatively predicted by this coarse-graining approach (yellow points, Fig. 4B). We conclude that treating Nar + Nir pairs as effective modules within larger communities recovers the predictive power of the additive consumer-resource model.

## Discussion

Quantifying the metabolic phenotypes of a diverse library of natural isolates using a consumerresource model allowed us to take a statistical approach to connecting genotypes to dynamical metabolic phenotypes. The outcome was a sparse mapping from gene content to single-strain metabolite dynamics that appears to exploit known mechanistic features of genes in the denitrification pathway to achieve predictive power. The resource-based modeling formalism permitted quantitative predictions of community-level metabolite dynamics. As a result, the approach yielded a direct mapping from genomic structure to metabolite dynamics at the community-level.

Our most surprising result is the extent to which gene presence and absence predicts metabolite dynamics at the strain and community-level. One might expect that the details of genomic or community context are necessary for predicting metabolite dynamics from genomes. Our results contradict this intuition, instead suggesting that, to a large extent, genes have a conserved impact on phenotypes irrespective of their genomic context. The result suggests that the observed conservation of gene content in wild communities has interpretable functional implications.

At the strain level, the apparent mechanistic relevance of the regression coefficients in this study suggests that a statistical approach, coupled with large-scale culturing and phenotyping on libraries of isolates (Connon and Giovannoni, 2002; Kehe et al., 2019), should be exploited to discover the salient features of genomes that determine diverse metabolic functions. Higher throughput measurements will enable a more detailed interrogation of genomic features, allowing us to extend our statistical approach to gene sequences and synteny. These insights might then be used to design genomes and communities with predefined metabolic capabilities by the addition or deletion of specific genes (Shaw et al., 2008). Data from phenotyping of diverse libraries of isolates could be combined with our approach to streamline the process of building constraint-based models for novel strains directly from genomic data (Norsigian et al., 2020) by allowing phenotypes to be predicted statistically rather than measured experimentally.

At the community level our approach could eventually enable the prediction of metabolite dynamics in complex communities where functional gene content has been assigned to individual genomes (Sieber et al., 2018). Soils and host-associated communities typically contain hundreds of bacterial taxa, so it may be necessary to test the predictive power of the consumer-resource formalism in communities of many taxa. However, micron-scale spatial structure in soils suggests that denitrification may occur locally, in communities of just a few taxa (Lensi et al., 1995), meaning that the genomic rules of denitrification discovered here are potentially relevant to important environmental contexts.

It is striking that communities containing both Nar and Nir phenotypes departed from the expectation of an additive model (Figure 3), and instead behaved as a single metabolic unit (Figure 4). We note that communities of Nar and Nir strains differ from a community containing Nar/Nir strains since they divide the denitrification cascade across two genomes. Consistent with our results (Figure 3A) and previous studies (Lilja and Johnson, 2016), segregating nitrate and nitrite reduction across genomes reduces the transient accumulation of nitrite during denitrification. Reducing transient nitrite accumulation is advantageous in low pH environments where nitrite is toxic (Lilja and Johnson, 2016). Our finding points to the possibility that Nar and Nir strains may have co-evolved to regulate nitrite accumulation, and this may reflect a community-level adaptation to acidic environments. Understanding the genomic control of the accumulation of intermediates during denitrification is essential for minimizing harmful nitrous oxide emissions (Cavigelli and Robertson, 2000) and controlling bacterial nitric oxide production in mammalian hosts (Hyde et al., 2014).

Understanding the evolutionary origins of the mapping from genomic structure to metabolic function is a key challenge going forward. Our study lends support to the idea that different genes in the denitrification pathway are adapted to different ecological niches (Jones and Hallin, 2010; Graf et al., 2014). In this light, our results potentially provide a route to understanding gene gain and loss statistics (Sela et al., 2019), or recent gene flow events (Arevalo et al., 2019), in terms of variation in metabolic phenotypes and their associated ecological niches. Uncovering the ecological niches and evolutionary forces that have shaped the sparse mapping from genomic structure to metabolic function could enable principled strategies for the design, control, and directed evolution of bacterial communities.

## Acknowledgements

We thank Laura Troyer for assistance with isolating bacterial strains, and Elizabeth Ujhelyi and Annette Wells for assistance with sequencing. We acknowledge Cameron Pittelkow for access to corn and soybean fields in Savoy, Illinois, and the laboratory of Julie Zilles for providing the bacterial strain *Paracoccus denitrificans* ATCC 19367. We also thank Rama Ranganathan, David Pincus, James Sethna, William Metcalf, Jun Song and members of the Kuehn laboratory and Mani group for helpful discussions. This work was supported by the National Science Foundation Division of Emerging Frontiers EF 2025293 (S.K.) and EF 2025521 (M.M.), the National Science Foundation Physics Frontiers Center Program PHY 0822613 and PHY 1430124 (S.K.), James S. McDonnell Foundation Postdoctoral Fellowship Award 220020499 (K.G.), and the Simons Foundation Investigator Award 597491 (M.M.).

## Author contributions

K.G.: Conceptualization, experimental design, data collection, formal analysis, coding, writing - original draft. D.P.: data collection. M.M.: Conceptualization, formal analysis, writing - revision & editing, supervision, funding acquisition. S.K.: Conceptualization, experimental design, formal analysis, writing - original draft, supervision, funding acquisition.

