## Supplementary material for "Genomic structure predicts metabolite dynamics in microbial communities": Methods and Supplemental Information

### Methods and Supplemental Information: Genomic structure predicts metabolite dynamics in microbial communities

**3 Supplemental Figures** **21**

**4 Supplemental Tables** **40**

**References** **49**

### 1 Methods

#### 1.1 Assay of nitrate and nitrite

Concentrations of nitrate and nitrite were measured following a modified version of the protocol by Miranda *et al.* (Miranda et al., 2001), where the conventional Griess assay for detection of nitrite is coupled with the chemical reduction of nitrate to nitrite using vanadium (III) chloride. Stock solutions of *N*-(1-Napthyl)ethylenediamine dihydrochloride (NEDD), sulfanilamide (SULF, in $5\%$ HCl), and $VCl_3$ (in $1\,\text{M}$ HCl) were prepared and stored as described in (Miranda et al., 2001). Griess reagent was freshly prepared on the day of each assay by mixing NEDD and SULF stock solutions with ultrapure $H_2O$ in a $5{:}5{:}9$ ratio.

Nitrate and nitrite were measured using $10\,\mu\text{L}$ samples of analyte in a 96-well microplate. Nitrite was first quantified by adding $190\,\mu\text{L}$ of Griess reagent to each sample and recording blank-corrected maximum absorbance within the interval $450$ to $650\,\text{nm}$, denoted $Abs^G_{max}$. Then nitrate was chemically reduced to nitrite by adding $50\,\mu\text{L}$ of $VCl_3$ solution to each sample and incubating at $30\,°\text{C}$ for $6$–$7\,\text{h}$, which was sufficient time for complete reduction of nitrate to nitrite (Fig. S1A). After incubation, the maximum blank-corrected absorbance within the interval $450$ to $650\,\text{nm}$, denoted $Abs^V_{max}$, was recorded to quantify the sum total of nitrate and nitrite in the sample.

Concentrations of nitrate and nitrite were determined using 8-point standard curves. Separate two-fold dilutions of $NaNO_2$ and $NaNO_3$ standards spanning $31\,\mu\text{M}$–$2\,\text{mM}$ were prepared, and $10\,\mu\text{L}$ samples of these standards were dispensed in triplicate into a 96-well plate, along with $10\,\mu\text{L}$ samples of ultrapure water that served as blanks. Griess reagents were used as described above to measure nitrite in the $NaNO_2$ standards, and these data were parameterized by fitting the equation,

$$Abs^G_{max}(I) = a_0 + a_1 I + a_2 I^2, \tag{1}$$

where $I$ is the concentration of $NO_2^-$. A quadratic term was included to account for the slight nonlinearity of the absorbance at high values of concentration. Then $VCl_3$ solution was used as described above to measure nitrate in the $NaNO_3$ standards, and these data were parameterized by fitting the equation,

$$Abs^V_{max}(A) = b_0 + b_1 A + b_2 A^2, \tag{2}$$

where $A$ is the concentration of $NO_3^-$. Example standard curves parameterized in this way are shown in Fig. S1B and C. Standard curves were run in triplicate on each day that an assay was performed.

The nitrate and nitrite concentrations in a new sample, for which $Abs^G_{max}$ and $Abs^V_{max}$ were recorded, were then determined by:

1. Solving equation (1) to determine $I$ from $Abs^G_{max}(I)$,

2. Solving equation (2) to determine the quantity $A+I$ from $Abs^V_{max}(A+I)$ (since this part of the assay measures the sum total of nitrate and nitrite in a sample),

3. Subtracting the result of step 1 from the result of step 2 to obtain $A$.

This approach was validated using mixed standards containing both nitrate and nitrite in a 1:1 ratio, and relative error was found to be typically less than $\pm5\%$ for concentrations $\geq 62.5\,\mu\text{M}$ (Fig. S1D).

In this study, nitrate and nitrite assays were performed directly on samples taken from denitrifying cultures (i.e., without first removing cells via filtration or centrifugation). To verify that the presence of cells does not significantly harm the quality of the measurement, error was evaluated in standards mixed with cell suspensions. An overnight culture of *Paracoccus denitrificans* ATCC 19367 was washed and diluted in phosphate buffered saline (PBS) to OD600 $\approx 0.4$ and then mixed in a 1:1 ratio with nitrate and nitrite standards. This produced samples with OD600 $\approx 0.2$, which is close to the maximum endpoint optical density observed in our denitrifying cultures grown with $2\,\text{mM}$ nitrate. In standards containing nitrate or nitrite separately, relative error of the assay is less than $\pm5\%$ for concentrations $\geq 125\,\mu\text{M}$, with relative error $\pm10\%$ at $62.5\,\mu\text{M}$ (Fig. S1E). In standards containing both nitrate and nitrite in a 1:1 ratio, relative error was again found to be typically less than $\pm5\%$ for concentrations $\geq 125\,\mu\text{M}$, with relative error $\pm10\%$ at $62.5\,\mu\text{M}$ (Fig. S1F). This indicates that the presence of cells increases measurement error only below concentrations of $125\,\mu\text{M}$.

#### 1.2 Isolation of denitrifying bacteria from soils

Denitrifying bacteria were isolated from soil samples following a modified version of the protocol developed by Lycus *et al.* (Lycus et al., 2017). $5$–$10\,\text{g}$ soil samples were collected from local prairie and forest environments, agricultural land, and manicured lawns (Table S3). Samples were stored separately in $50\,\text{mL}$ centrifuge tubes at $4\,^{\circ}\text{C}$ for no longer than three months before use.

Each soil sample was prepared for isolation by combining with $25\,\text{mL}$ PBS (pH 7.4) and $5$–$10\,\text{g}$ sterile $4\,\text{mm}$ glass beads. Additionally $0.5\,\text{mL}$ cycloheximide solution ($10\,\text{mg/mL}$) and $20\,\mu\text{L}$ nystatin solution ($25\,\text{mg/mL}$) were added to prevent fungal growth. In certain rounds of isolation (Table S3), sterile $NaNO_2$ solution ($1\,\text{M}$) was added to enrich for nitrite-reducing bacteria. Tubes were then vortexed (Vortex-Genie 2) at high speed for $1\,\text{min}$ to homogenize the samples. In certain rounds of isolation (Table S3), homogenized soils were incubated at either room temperature or $30\,^{\circ}\text{C}$ for up to two weeks, and then briefly re-vortexed before further processing.

Supernatants from homogenized soils were then diluted and plated. After vortexing, large particles in homogenized soil samples were allowed to settle for $20$–$30\,\text{min}$ before transferring $1\,\text{mL}$ of supernatant to sterile $1.7\,\text{mL}$ microcentrifuge tubes. Soil supernatants were then serially-diluted in PBS to obtain $10^{-4}$-fold and $10^{-5}$-fold dilutions. $100\,\mu\text{L}$ of the $10^{-4}$ and $10^{-5}$ soil supernatant dilutions were plated on 1/10X tryptic soy agar (1/10X TSA, $1.5\,\text{g/L}$ tryptone, $0.5\,\text{g/L}$ soytone, $0.5\,\text{g/L}$ NaCl, $15\,\text{g/L}$ agar), with two replicates for each dilution. Plates were then incubated under aerobic conditions at $30\,^{\circ}\text{C}$ for $48\,\text{h}$.

Colonies from plated soil supernatants were picked and streaked to purity. Plated soil dilutions were examined for growth after incubation. Plates showing little or no growth were incubated for an additional 24–48 h until colonies appeared. Plates showing likely fungal growth were discarded. For each set of plates derived from a soil sample, 5–15 well-separated colonies were picked and streaked to purity on 1/10X TSA plates, again incubating at 30 °C. Whenever possible, colonies were selected which varied in morphology, size, and color, in order to enhance the diversity of the isolate collection.

Bacterial isolates were assayed for nitrate and/or nitrite-reduction capability. Isolates were first cultured aerobically in sterile 96-deepwell plates (Axygen PDW20C) containing 1 mL per well 1/10X tryptic soy broth liquid medium (1/10X TSB), inoculated directly from streak plate colonies. Deepwell plates were sealed with a gas-permeable membrane (Diversified Biotech BERM-2000) and incubated/shaken at 30 °C/950 RPM (Talboys Professional 1000MP) for 48 h. 1 µL of each dense aerobic cultures was then passaged to fresh 96-deepwell plates containing 1 mL per well 1/10X TSB supplemented with either sterile $NaNO_3$ or $NaNO_2$ solution (1 M) to yield a final concentration of 2 mM $NO_3^-$ or $NO_2^-$, with two replicates of each condition per isolate. Plates were sealed with gas-permeable membranes (Diversified Biotech BEM-1) and transferred to an $N_2$-purged anaerobic glove box (Coy Laboratory Products 7601-110/220). Plates were incubated/shaken at 30 °C/950 RPM for 48 h under anaerobic conditions. After incubation, plates were sampled to determine endpoint concentrations of nitrate and nitrite.

Nitrate and nitrite-reduction capability for each isolate was determined by comparing endpoint concentrations of $NO_3^-$ and $NO_2^-$ in each culture with the corresponding concentrations in uninoculated controls. Isolates showing lower endpoint levels of nitrate and/or nitrite than the uninoculated controls were identified as nitrate and/or nitrite reducers, respectively. Strains that performed both nitrate and nitrite reduction were classified as “Nar/Nir” strains, while strains that performed nitrate or nitrite reduction only were classified as “Nar” or “Nir” strains, respectively. These isolates were cryopreserved by mixing 500 µL of saturated culture grown in 1/10X TSB under aerobic conditions with 500 µL filter-sterilized 50% glycerol solution in 2 mL cryotubes, and freezing/storing at −80 °C.

#### 1.3 Defined growth medium

In order to eliminate the possibility of fermentative metabolism in subsequent (post-isolation) denitrification experiments, a defined growth medium designed by Heylen *et al.* (Heylen et al., 2006) for the cultivation of diverse denitrifying bacteria was used (Table S4). The medium contained 25 mM succinate as the sole (non-fermentable) carbon source, 15 mM ammonium as the assimilatory nitrogen source, and a 40 mM phosphate buffer with the final medium pH adjusted to 7.3. The medium additionally contained trace metals and vitamins. In denitrification experiments using this medium, nitrate or nitrite was provided in at most 2 mM concentrations, and it was expected that carbon, ammonia, and phosphate were in relative excess, so that nitrate or nitrite was the growth-limiting nutrient. It was expected that nitrate and nitrite assimilation were inhibited by high concentrations of ammonia (Zumft, 1997).

Denitrifying isolates were screened for growth on succinate defined medium (SDM) under aerobic conditions by first growing axenically from freezer stocks in 1/10X TSB, passaging 1:300 into 300 µL SDM in 96-well plates (with two replicates per isolate), and culturing at 30 °C for 48 h. Endpoint optical density was measured to assess growth on SDM. Denitrifying isolates unable to grow on SDM (approximately 36% of isolates) were excluded from further experimentation and analysis.

#### 1.4 Denitrifying conditions

Post-isolation denitrification experiments were performed in a vinyl glove box (Coy Laboratory Products 7601-110/220) purged of oxygen with a 99%/1% $N_2$/$CO_2$ gas mixture. Provision of $CO_2$ was necessary to support the growth of cultures from low initial cell densities (OD600 $\ll 0.01$), likely due to the $CO_2$-fixation requirements of core anaplerotic metabolism (White et al., 2012). Gas mixing and a purge rate of 20 SLPM were controlled using digital mass flow controllers (Sierra Instruments SmartTrak 50). Oxygen and $CO_2$ concentrations inside the glove box were continually monitored using Arudino-attached (Mathupala et al., 2016) optical sensors (SST Sensing LOX-02, Gas Sensing Solutions EXPLORIR-M-20). Gaseous oxygen concentration was maintained below 50 ppm, which was sufficient to prevent growth via aerobic respiration. This was verified using two denitrifying isolates, the Nar strain *Paracoccus* sp. PAR01 and the Nir strain *Pseudomonas* sp. PDM13, which were grown under denitrifying conditions for 64 h with and without a suitable electron acceptor (i.e., nitrate for the Nar strain and nitrite for the Nir strain). The Nar strain PAR01 was inoculated at cell density $\text{OD600}_0 = 0.011$ and grew to $\text{OD600}_{end} = 0.072$ with nitrate and remained at $\text{OD600}_{end} = 0.013$ without nitrate. Similarly, the Nir strain PDM13 was inoculated at cell density $\text{OD600}_0 = 0.010$ and grew to $\text{OD600}_{end} = 0.026$ with nitrite and remained at $\text{OD600}_{end} = 0.010$ without nitrite.

#### 1.5 Denitrification dynamics experiments

Denitrifying strains were pre-cultured under aerobic conditions (using first broth then SDM) prior to growth under denitrifying (anaerobic) conditions in SDM. Strains were first grown axenically to saturation from freezer stocks in 1/5X TSB, then passaged 1:100 into SDM and again grown to saturation. All pre-cultures were grown in 24-well plates containing 1.7 mL medium per well and incubated/shaken at 30 °C/400 RPM under aerobic conditions. Since strains in our collection vary widely in the time required to grow to saturation in 1/5X TSB and SDM (12–80 h), inoculation and passaging of strains was timed so that SDM aerobic cultures would enter stationary phase within 12 h of the start of an experiment in denitrifying conditions (Table S5).

Experiments to measure the nitrate/nitrite-reduction dynamics of strains in monoculture were performed at multiple initial media and cell density conditions: (a) 2 mM $NO_*^-$, $\text{OD600}_0$ = 0.01, (b) 1 mM $NO_*^-$, $\text{OD600}_0$ = 0.01, (c) 2 mM $NO_*^-$, $\text{OD600}_0$ = 0.001, where Nar/Nir strains were cultured with $NO_3^-$ and $NO_2^-$ separately (6 conditions total), Nar strains were cultured with

$NO_3$ (3 conditions total), and Nir strains were cultured with $NO_2^-$ (3 conditions total). SDM containing (separately) $2\,\text{mM}$ $NO_3^-$, $1\,\text{mM}$ $NO_3^-$, $2\,\text{mM}$ $NO_2^-$, and $1\,\text{mM}$ $NO_2^-$ were prepared by supplementing fresh SDM with sterile $NaNO_3$ or $NaNO_2$ ($1\,\text{M}$) solutions. Aerobic SDM pre-cultures were normalized to two levels of optical density (OD600 $\approx 1.2$ and $0.12$) by diluting with PBS, and these normalized densities were recorded. $96$-deepwell plates containing $1.2\,\text{mL}$ of $NO_3^-$ or $NO_2^-$-supplemented SDM were then inoculated with $10\,\mu\text{L}$ of density-normalized culture, resulting in OD600 $\approx 0.01$ and $0.001$ cell density conditions. $4$ replicates were used for each combination of media and cell density conditions.

Inoculated plates were sealed with gas-permeable membranes (Diversified Biotech BERM-2000) and transferred to an anaerobic glove box for incubation/shaking at $30\,°\text{C}/950\,\text{RPM}$. Cultures were manually sampled with a multichannel pipette ($10\,\mu\text{L}$ per well) at 2, $4$, $6$, $8$, $16$, $32$, and $64\,\text{h}$ from the start of anaerobic culture. Samples were stored in clean $96$-well plates, and immediately sealed (Bio-Rad MSB1001) and frozen at $-20\,°\text{C}$ for later assay of nitrate and nitrite. Additionally, at $64\,\text{h}$, $300\,\mu\text{L}$ per well of culture was sampled into clean $96$-well plates and the optical density (pathlength normalized to $1\,\text{cm}$) immediately measured to quantify endpoint abundances. Strains observed to form excessive cell aggregation (to the extent that quantification of endpoint abundances by optical density was not possible) were at this point excluded from further experimentation and analysis.

Additional experiments to measure nitrate/nitrite-reduction dynamics on communities of 2–3 strains (including monoculture controls) were performed in separate $2\,\text{mM}$ $NO_3^-$ and $2\,\text{mM}$ $NO_2^-$ conditions. Aerobic pre-cultures were normalized to OD600 $\approx 1.2$ with PBS. Using liquid handling robotics (Formulatrix Mantis), $96$-deepwell plates containing $1.2\,\text{mL}$ of $2\,\text{mM}$ $NO_3^-$ or $NO_2^-$ SDM was inoculated with $10\,\mu\text{L}$ droplets of density-normalized culture in defined combinations (e.g., a pair community containing strains A and B would be inoculated with separate $10\,\mu\text{L}$ droplets of strain A and strain B). Following inoculation, all culture and sampling details were the same as described above.

Measurements were post-processed to correct for gradual evaporation of $H_2O$ that increased apparent concentrations with time. $3$–$8$ uninoculated controls containing nitrate or nitrite-supplemented medium were used on each culture plate to quantify the effect of evaporation (Fig. S2A). Raw nitrate and nitrite measurements for cultures were scaled at each time point by the concentration of initially supplied nitrate or nitrite divided by the median concentration across blank measurements at that time point (see example in Fig. S2B and C).

#### 1.6 Whole genome sequencing and annotation

Denitrifying strains were grown axenically from freezer stocks or plated colonies in 1/10X TSB incubated at $30\,°\text{C}$ with shaking. Saturated cell cultures were harvested and DNA extracted using the DNeasy UltraClean Microbial Kit (Qiagen). DNA concentrations were quantified using the Qubit dsDNA BR Assay Kit (Invitrogen).

Library preparation for sequencing of DNA extracts was performed using the Nextera DNA Flex Library Prep Kit (Illumina). Barcoded libraries for each strain were pooled in groups of 21–24 strains and quantified using the Qubit dsDNA BR kit and an Agilent 2100 Bioanalyzer (Carver Biotechnology Center, University of Illinois at Urbana-Champaign). Each pooled library was then separately sequenced using a MiSeq Reagent Kit v3 (Illumina, $2 \times 300\,\mathrm{bp}$ paired-end), with a $12\,\mathrm{pM}$ library loading concentration and a $1\%$ spike-in of PhiX Control v3 (Illumina). Sequencing was performed on a locally maintained and operated Illumina MiSeq system.

Raw paired-end reads were trimmed of low-quality regions and Illumina adapters using Trimmomatic 0.39 (Bolger et al., 2014). Trimmed reads were assembled into contigs *de novo* using SPAdes 3.13.0 (Bankevich et al., 2012) with k-mer lengths of 21, 33, 55, 77, 99, and 127 and the error-correction option "careful" enabled. Assembly quality was assessed using QUAST 5.02 (Gurevich et al., 2013). Our draft assembly of the laboratory strain *Paracoccus denitrificans* ATCC 19367 was compared to a complete assembly of the same strain by Si *et al.* (Si et al., 2019), which served as a reference. The comparison indicated a $97.8\%$ genome fraction (percentage of bases in the reference that align to our assembly), with one $30.8\,\mathrm{kbp}$ relocation missassembly, $7.02$ mismatches (sequencing errors or single nucleotide polymorphisms) per $100\,\mathrm{kbp}$, and $0.43$ indels per $100\,\mathrm{kbp}$. Our assembly produced $4635$ predicted genes relative to $4644$ genes predicted in the reference. The coverage depth of our *P. denitrificans* draft assembly was $32$X, compared to a median of $35$X for the other strains in our collection. Together these analyses suggest that our draft assemblies are likely to cover the vast majority of the protein-coding sequences, with relatively few errors that would affect the inference of gene presence or absence.

Contigs were uploaded for gene annotation on the RAST Server (http://rast.theseed.org) using the RASTtk pipeline (Brettin et al., 2015). Denitrification gene presence/absence information was obtained from annotation files by a text search of gene function labels (Table S6). The *NarXL* two-component nitrate/nitrite sensing system was considered "present" if a gene encoding the sensor (*narX*) and/or the response regulator (*narL*) were identified in the annotation. Each gene identified as a *NarK*-type nitrate transporter was classified post-annotation as encoding either a nitrate/$H^+$ symporter (*narK1*), a nitrate/nitrite antiporter (*narK2*), or a fusion of both transporters (*narK1K2*). This classification was performed by locally aligning each transporter sequence with the *narK1K2* gene previously identified in *Paracoccus denitrificans* PD1222 (Goddard et al., 2017), for which the N-terminal domain has been identified to be *NarK1* and the C-terminal domain *NarK2*. From this it was determined which RAST gene function labels correspond to *narK1*, *narK2*, and *narK1K2*.

Additionally annotation files were searched for the cytochrome c nitrite reductase gene (*nrfA*) associated with dissimilatory nitrate reduction to ammonia (DNRA). Strains possessing this gene were excluded from further experimentation and analysis.

#### 1.7 Phylogenetic classification of denitrifying strains

Phylogenetic analysis of denitrifying strains was performed using full 16S rRNA sequences identified in annotated draft genome assemblies. 16S sequences were uploaded for classification up to the genus level using the SILVA ACT service (http://www.arb-silva.de/) (Pruesse et al., 2012). A maximum-likelihood phylogenetic tree was computed using MEGA X 10.1.8 (Kumar et al., 2018).

### 2 Supplemental Information

#### 2.1 Consumer-resource model for nitrate and nitrite dynamics

We formulated a consumer-resource model to parameterize the dynamics of nitrate and nitrite-reduction. For a single strain that performs both nitrate and nitrite reduction (Nar/Nir strain), the model is as follows:

$$\begin{aligned} \frac{dx}{dt} &= \left(\gamma_A r_A \frac{A}{K_A + A} + \gamma_I r_I \frac{I}{K_I + I}\right) x \\ \frac{dA}{dt} &= -r_A \frac{A}{K_A + A} x \\ \frac{dI}{dt} &= \left(r_A \frac{A}{K_A + A} - r_I \frac{I}{K_I + I}\right) x. \end{aligned} \tag{3}$$

The variable $x$ is the cell density of a single population, and $A$ and $I$ are the concentrations of nitrate and nitrite respectively. The parameters $\gamma_A$ and $\gamma_I$ are the growth yields, $r_A$ and $r_I$ are the rates of reduction per unit cell density, and $K_A$ and $K_I$ are the substrate affinity constants for nitrate and nitrite respectively.

The model, which resembles the Monod model for bacterial growth (Monod, 1949), assumes that growth of the population occurs at a rate proportional to the reduction rates of nitrate and nitrite, with the two resources treated as substitutable. Nitrite is the direct product of nitrate reduction (in a 1:1 stoichiometry). Nitric oxide, the product of nitrite reduction, is not explicitly modeled, and thus mass leaves the system when nitrite is reduced. For a strain that performs only nitrate reduction (Nar strain), the model simplifies by setting $r_I = 0$. Likewise for a strain that performs only nitrite reduction (Nir strain), $r_A = 0$.

#### 2.2 Inferring consumer-resourcing parameters from data

We cultured 78 soil isolates and one reference strain *Paracoccus denitrificans* ATCC 19367 in monoculture under denitrifying conditions (see §1.5 for experimental details) in order to estimate parameters to the consumer-resource model (3). We then removed strains with low quality fits, yielding a final strain library of 62 strains.

##### 2.2.1 Fitting yield parameters

We directly measured the yield parameters $\gamma_A$ and $\gamma_I$ using endpoint measurements of cell abundance and nitrate/nitrite concentration. Strains were grown in multiple initial conditions:

(a) $2\,\mathrm{mM}$ $NO_*^-$, $OD600_0$ = $0.01$, (b) $1\,\mathrm{mM}$ $NO_*^-$, $OD600_0$ = $0.01$, (c) $2\,\mathrm{mM}$ $NO_*^-$, $OD600_0$ = $0.001$, where Nar/Nir strains were cultured with $NO_3^-$ and $NO_2^-$ separately ($6$ conditions total), Nar strains were cultured with $NO_3$ ($3$ conditions total), and Nir strains were cultured with $NO_2^-$ ($3$ conditions total). In all conditions, it was expected that nitrate and/or nitrite were the growth-limiting nutrients. Yield coefficients were then obtained by fitting the equation

$$\begin{aligned} \Delta\mathrm{OD600} = \mathrm{OD600}_{end} - \mathrm{OD600}_0 &= \gamma_0 + \gamma_A \Delta A + \gamma_I \Delta I \\ &= \gamma_0 + \gamma_A (A_0 - A_{end}) + \gamma_I (A_0 - A_{end} + I_0 - I_{end}), \end{aligned} \tag{4}$$

where $\mathrm{OD600}_{end}$ is the endpoint optical density measurement, $\mathrm{OD600}_0$ is the initially prescribed optical density, $A_0$ and $I_0$ are initially prescribed concentrations of nitrate and nitrite respectively, and $A_{end}$ and $I_{end}$ are endpoint measurements of nitrate and nitrite respectively. The $\Delta A$ and $\Delta I$ terms represent total nitrate and nitrite reduced, respectively. The quantity $A_0 - A_{end}$ is included in the $\Delta I$ term because nitrite is the product of nitrate reduction. The intercept term $\gamma_0$ is typically close to zero (Fig. S3A). For each strain, this equation is fit to all initial cell density and media conditions combined using ordinary least squares (see examples in Fig. S3B to D). Optical densities were converted into nominal cell densities assuming OD600 = $0.1$ corresponds to $10^8$ $\mathrm{cells/mL}$ (BioNumbers ID 100985) to give yields in units of $\mathrm{mmol}^{-1}$.

##### 2.2.2 Fitting rate parameters

For each strain, having determined the yield parameters, we then globally (simultaneously) fit the rate parameters $r_A$ and $r_I$ to nitrate/nitrite dynamics data across all experimental conditions by minimizing the sum of squared residuals between the data and numerical solutions to equation (3) (see examples in Fig. S4). We solved equation (3) numerically using the differential equation solver ode23s in MATLAB R2017b. We initialized the solver at $t = t_1$ (where $t_1$ is the time point of first nitrate/nitrite measurement), taking initial conditions $N(t_1) = \mathrm{OD600}_0$, and $A(t_1)$ and $I(t_1)$ to the median measured values of nitrate and nitrite concentration at $t = t_1$ over experimental replicates within a given condition. We used the constrained optimization function fmincon in conjunction with the global minimum search function GlobalSearch to minimize the sum of squared residuals and obtain optimal values of $r_A$ and $r_I$. Values of $r_A$ and $r_I$ were constrained between $0$ and $5 \times 10^{-8}\,\mathrm{mmol/h}$.

Initially we used this approach to simultaneously fit both rates ($r_A$ and $r_I$) and substrate affinities ($K_A$ and $K_I$), but found in nonparametric bootstrap estimates of parameter error (described later in §2.2.4) that the affinities $K_A$ and $K_I$ are not well constrained by our experimental measurements (Fig. S5A). This is likely because the true values are small relative to the typical scale of substrate concentrations in our experiments; values for nitrate/nitrite affinity constants between $0.003$–$0.055\,\mathrm{mM}$ have been identified for denitrifying cultures previously (Beccari et al., 1983; Claus and Kutzner, 1985; Kornaros et al., 1996; Dinçer and Kargı, 2000), while we initialized cultures with $1$–$2\,\mathrm{mM}$ nitrate/nitrite. We remark that in the dynamical regimes of equation (3) where substrate concentrations are much greater than affinity constants (i.e., $A \gg K_A$, $I \gg K_I$), the dependence of (3) on $K_A$ and $K_I$ vanishes. This restricts identifiability of the affinity parameters to the regime where $A, I$ are on the same scale as $K_A, K_I$ (likely in the micromolar range). This regime typically occurs only briefly during our experiments (just before the substrates are exhausted), and sampling this regime adequately would require a much greater sampling frequency than what we have attempted in this study. Since qualitative behavior (Fig. S5B) and fit quality (Fig. S5C) are insensitive to $K_A$ and $K_I$, we fixed $K_A = K_I = 0.01\,\text{mM}$.

##### 2.2.3 Excluding poorly fitting strains

We observed two categories of strains with poor model fits that were excluded from further experimentation and analysis. First we observed 7 Nar strains for which nitrate concentrations appear to asymptotically approach non-zero values (see example in Fig. S6A). This behavior is not permitted by the model (3), since the steady state value of $A$ must be $0$ whenever $r_A, x > 0$. Second, we observed $10$ strains that have poor model fits evidenced by relatively large values of root-mean-square errors (RMSE) evaluated between observed data and model solutions across all experimental conditions. A threshold value of RMSE $= 0.17$ separated these poorly fitting strains from the others. For these strains we typically observed that global fits did well at fitting some experimental conditions and poorly at others (e.g., fitting the data well for conditions where nitrite is initially supplied and fitting poorly when nitrate is initially supplied, as in the example shown in Fig. S6B), again indicating an inconsistency with the modeling framework (3). We concluded in both cases that appropriately capturing dynamics requires additional mechanisms not currently contained in (3) and excluded these strains from the final library.

##### 2.2.4 Estimating parameter error

We used a nonparametric bootstrap to estimate the sampling distributions of $\{r_A, \gamma_A, r_I, \gamma_I\}$ for the remaining $62$ strains. We generated bootstrap datasets by resampling (with replacement) among replicates; that is, for an experimental condition (e.g., $\text{OD600}_0 = 0.01$, $A_0 = 2\,\text{mM}$, $I_0 = 0\,\text{mM}$) performed with four replicates, we created a new dataset by randomly selecting a set of four replicates with replacement, doing this for all conditions. We then used the approach described above to re-fit $\{r_A, r_I, \gamma_A, \gamma_I\}$ to these bootstrap resamples. We repeated this $100$ times for each strain to obtain sampling distributions for these parameters. The 25th to 75th percentiles of the sampling distributions determined using this approach are shown in Fig. S7A. Fractional errors, defined as the ratio of the interquartile range to the value of the parameter, are less than $10\%$ for the vast majority of inferred parameters (Fig. S7B). Since $\gamma_A$ and $\gamma_I$ inferences are error-prone when measured cell abundances (and therefore yields) are low, we set a yield parameter identically to zero if its estimate is negative or within one standard error of zero.

#### 2.3 Regressing consumer-resource parameters onto denitrification gene presence and absence

##### 2.3.1 Formulating the regression problem

We used linear regression to predict the measured consumer-resource model parameters $\{r_A, \gamma_A, r_I, \gamma_I\}$ from the presence/absence of denitrification-related genes in the genomes of each strain. We formulated the regression problem as follows:

$$y_i = \beta_0 + \sum_{j=1}^{P} \beta_j g_{ij} + \varepsilon_i \quad \text{for } i = 1, \ldots, N. \tag{5}$$

The response variable $y_i$ is the observed value of a consumer-resource model parameter (e.g., $r_A$) for strain $i$, with $N$ such observations in total. The predictors $g_{ij}$ are indicator variables taking the value 1 if strain $i$ has gene $j$ and taking zero otherwise, with $P = 17$ predictors in total. The regression coefficients $\beta_j$ and intercept $\beta_0$ are fit by the regression, which determines the residual term $\varepsilon_i$. This formulation assumes that genes have independent and additive effects on the value of a phenotypic parameter.

We fit equations of this form separately for each consumer-resource model parameter, obtaining different $\vec{\beta}$ and $\beta_0$ for each regression. Note that for nitrate-related parameters ($r_A$ and $\gamma_A$), only strains capable of nitrate reduction (Nar and Nar/Nir phenotypes) were included in the predictor-response datasets ($N = 58$ strains). Similarly for the nitrite-related parameters, only strains with Nar/Nir and Nir phenotypes were included in the predictor-response datasets ($N = 47$ strains). We also note that prior to fitting, all predictors were standardized to have zero mean and unit variance.

We used the LASSO regression method (Hastie, R. Tibshirani, and Friedman, 2008; Hastie, R. Tibshirani, and Wainwright, 2016) in MATLAB R2017b to solve (5). For a given predictor matrix $X$ and response vector $y$, LASSO regression solves

$$\min_{\beta_0, \vec{\beta}} \left\{ \frac{1}{2N} \sum_{i=1}^{N} \left( y_i - \beta_0 - x_i^T \vec{\beta} \right)^2 + \lambda \|\vec{\beta}\|_1 \right\}, \tag{6}$$

performing both variable selection and regularization by penalizing the sum of squared residuals by the $L_1$ norm of the coefficient vector $\vec{\beta}$. The strength of the penalty is controlled by the hyperparameter $\lambda$, which at moderate values sets the coefficients of poor predictors identically to zero, thus resulting in a sparse model. Typically the hyperparameter value $\lambda = \hat{\lambda}$ is selected by minimizing prediction error in cross-validation, which optimizes the ability of the model to generalize out of sample, and makes the method suitable for datasets where overfitting via an approach such as ordinary least squares (OLS) is likely because the number of predictors and the number of observations are on the same order of magnitude.

##### 2.3.2 Using cross-validation to determine optimal hyperparameter values

For each regression of the form given in equation (5), we used iterated $K$-fold cross-validation to determine the value $\lambda = \hat{\lambda}$ that minimizes estimated prediction error. Let $\Lambda$ be a set of discrete values of $\lambda$ for which equation (6) is solved. Each iteration of cross-validation proceeds in the following way:

1. Randomly partition the data into $K$ subsamples of roughly equal size.
2. For each $k = 1, 2, ..., K$:
   (a) Holding out the $k$th fold (test set), solve (6) at each $\lambda \in \Lambda$ using the union of the remaining remaining $K - 1$ folds of data (training set).
   (b) Use the resulting regression coefficients to predict the responses in the test set.
   (c) Evaluate root-mean-square error of the test set prediction.

This process is iterated $M$ times, each time choosing a new random partition of the data into $K$ subsamples. We took the average of the $KM$ estimates of prediction error at each value of $\lambda \in \Lambda$, and the optimal value $\lambda = \hat{\lambda}$ was then selected as that which minimized this estimate of error. We chose an iterated $K$-fold cross-validation approach over either basic $K$-fold or leave-one-out ($K = N$) cross-validation because iterated cross-validation averages over the sampling variability inherent in randomly partitioning the dataset into groups, and also because partitioning into $K \neq N$ folds allows us to estimate the out-of-sample performance of the model through statistics such as the coefficient of determination between observed and predicted values on each test set (see §2.3.3).

The values of $\hat{\lambda}$ selected via iterated 4-fold cross-validation ($M = 10^4$ iterations) are shown in Fig. S8A. We chose $K = 4$ to balance the relative sizes of training and test sets for the purpose of evaluating out-of-sample performance; as $K$ increases, the size of the cross-validation training set increases, while the size of test set decreases. We determined that $\hat{\lambda}$ was insensitive to the choice of $K$ (Fig. S8B).

##### 2.3.3 Using cross-validation to assess out-of-sample performance

Out-of-sample performance of regressions on each consumer-resource parameter was estimated during iterated $K$-fold cross-validation (described in §2.3.2) by computing the coefficient of determination ($R^2$) between observed and predicted response values in each test set. The distributions of these values obtained at $\lambda = \hat{\lambda}$ for iterated 4-fold cross-validation ($M = 10^4$ iterations), denoted $R^2_{CV}$, are shown in Fig. S9A. The median of each distribution, denoted $\bar{R}^2_{CV}$, is positive for all regressions.

The significance of $\bar{R}^2_{CV}$ for regressions on each consumer-resource parameter was evaluated under the null hypothesis that there is no relationship between the predictor and the response via a permutation test. The response variable for each regression was repeatedly permuted ($10^3$ permutations) and regression coefficients re-fitted, using iterated 4-fold cross-validation ($M = 10^2$ iterations) to select the hyperparameter for each permutation. The null distributions of $\bar{R}^2_{CV}$ computed using this approach are shown in Fig. S9B, with $p < 0.001$ in all cases.

###### 2.3.4 Post-selection inference

We performed post-selection inference on regression coefficients using the package selectiveInference 1.2.5 in R 3.6.1. When the data used for model training are also used for inference, it is necessary to account for the fact that LASSO performs variable selection (Hastie, R. Tibshirani, and Wainwright, 2016; Taylor and R.J. Tibshirani, 2015). Since LASSO selects variables with high predictive power in the training set, neglecting variable selection results in overoptimistic confidence intervals and $p$-values. We used the function fixedLassoInf in the selectiveInference package to obtain $p$-values for each nonzero regression coefficient under the null hypothesis the true value is zero, conditional on the fact the variable was selected by LASSO to have a nonzero coefficient at $\lambda = \hat{\lambda}$ (Fig. S10).

###### 2.3.5 Investigating the impact of phylogenetic correlation in the strain library

While closely related strains in our library displayed variability in both the phenotypic parameters $\{r_A, \gamma_A, r_I, \gamma_I\}$ (Fig. 1) and in denitrification gene presence/absence (Fig. 2), we sought to determine whether possible phylogenetic correlations (Martiny et al., 2015) between traits and genes influenced either regression performance or the coefficients selected by LASSO. To do this, we effectively "collapsed" clades comprising strains with identical 16S rRNA sequences (i.e., the strains that exhibit the greatest degree of phylogenetic correlation) by randomly selecting one representative and removing the remaining strains from the dataset. This removed $12$ strains from the library, resulting in a subsampled dataset of $50$ strains (Fig. S11A). We then performed LASSO regressions on this subsampled dataset as described in §2.3.1-2.3.3. The results of these regressions are shown in Fig. S11B. We observed that both in-sample and out-of-sample metrics of model performance ($R^2_{fit}$ and $\bar{R}^2_{CV}$, respectively) changed little relative to the corresponding values for regressions on the full dataset (cf. Fig. 2C to F). Similarly, we observed that the regression coefficients did not change substantially (cf. Fig. 2G to J). We concluded therefore that phylogenetic correlations did not substantially impact regressions on our full dataset.

##### 2.4 Comparing denitrification genes with randomly-selected genes as predictors for consumer-resource parameters

In order to assess whether denitrification genes are better predictor variables than other possible sets of genes, we regressed the consumer-resource parameters $\{r_A, \gamma_A, r_I, \gamma_I\}$ onto the presence/absence of randomly-selected genes from the set of all annotated genes in strains in our library.

We first identified the set of all uniquely-labelled protein-encoding genes present in the RAST annotations for the $62$-strain library. From this set of $12\,864$ unique genes, we removed $50$ genes associated with denitrification, removing not only terminal reductases, sensors/regulators, and transporters that were included as variables in the regression problem (5) (Table S6), but also related chaperones, structural genes, and biosynthesis genes that frequently occur in clusters with the denitrification genes (Table S7).

We then generated new sets of predictor variables by randomly selecting genes from the set of $12\,814$ non-denitrification genes and constructing binary presence/absence matrices as we did for denitrification-related genes. In order to create a fair comparison between these randomly-selected predictors and the denitrification gene predictors, we (a) selected sets of only 17 genes, matching the dimensionality of the denitrification gene presence/absence matrix, and (b) performed rejection sampling to match the gene presence frequency (fraction of strains that possess a given gene) distribution of the denitrification genes. The latter was a necessary consideration because the distribution of presence frequencies of non-denitrification genes is heavily skewed toward zero relative to the frequencies of denitrification genes (Fig. S12A), indicating that a large portion of genes occur only in a small number of strains.

To perform rejection sampling, we estimated densities for the gene frequency distributions and used these densities to define acceptance probabilities for random samples. Let $f(x)$ denote the probability density of gene frequencies for non-denitrification genes and similarly $g(x)$ for denitrification genes, where $x$ here denotes gene presence frequency, and set the constant $H$ so that $g(x)/f(x)/H \leq 1$ for all $x \in [0, 1]$. We then performed rejection sampling in the following way:

1. With uniform probability, randomly sample a candidate gene from the set of non-denitrification genes. Denote the frequency of this gene as $x_0$.

2. Accept the candidate gene as a predictor with probability $g(x_0)/f(x_0)/H$. Otherwise reject the candidate and return to step 1.

3. Return to step 1 until 17 genes are accepted.

This process results in a set of 17 predictors that have a gene presence frequency distribution approximately equal to $g(x)$. We used the ksdensity function in MATLAB R2017b to estimate $f(x)$ and $g(x)$ using a bandwidth parameter value of $0.4$ (Fig. S12A). We used this rejection sampling approach to generate $10^3$ different sets of 17 predictors for regressing against each of the phenotypic parameters $\{r_A, \gamma_A, r_I, \gamma_I\}$.

We performed LASSO regressions on each of the consumer-resource parameters $\{r_A, \gamma_A, r_I, \gamma_I\}$ using the rejection-sampled sets of gene predictors, where the optimal hyperparameter value $\lambda = \hat{\lambda}$ was selected via iterated $4$-fold cross-validation ($M = 10^3$ iterations) (see §2.3.2 for details). Out-of-sample performance of each regression was also evaluated in cross-validation by computing $\bar{R}^2_{CV}$, the median coefficient of determination value computed across cross-validation test sets (see §2.3.3 for details). The distributions of these $\bar{R}^2_{CV}$ values across different sets of random genes are shown in (Fig. S12B), alongside the values of $\bar{R}^2_{CV}$ obtained via the regressions onto denitrification-related genes. We observed that the denitrification genes perform better than the typical set of random genes in regressions on three out of four phenotypic parameters. On the whole, this analysis suggests that denitrification genes are better predictors than arbitrary genes, but it remains unclear why arbitrary genes tend have modest predictive power ($\bar{R}^2_{CV} > 0$), and why the denitrification genes perform worse than the typical set of random genes for the $r_I$ parameter. One possible explanation is that some random genes serve as good predictors only because they resemble the denitrification genes in terms of presence and absence.

We therefore asked whether the predictive power of random genes arises because of correlations (in terms of presence/absence) with the denitrification genes. To answer this question, we constructed new sets of random gene predictors via rejection sampling as described above, but this time sampling only from the set of genes that have low or insignificant correlation with denitrification genes. Specifically, for each of the non-denitrification gene presence/absence vectors, we computed the Pearson correlation $\rho$ with each of the denitrification gene presence/absence vectors. We then identified the set of non-denitrification genes that have large correlations ($|\rho| \geq 0.5$) with any of the denitrification genes that are significant at the $1\%$ level, where we evaluated significance by generating a zero-correlation null distribution via a permutation test. The latter consideration is important for evaluating correlations between very high or very low-frequency genes, for which the probability of spurious large correlations can be high. We excluded these denitrification gene-correlates from sampling when generating the new sets of random gene predictors. As before, we generated $10^3$ different sets of 17 predictors for regressing against each of the consumer-resource parameters.

We again performed LASSO regressions on each of the consumer-resource parameters $\{r_A, \gamma_A, r_I, \gamma_I\}$ using the rejection sampled genes that exclude denitrification gene correlates. The distributions of the resulting $\bar{R}^2_{CV}$ values are shown in (Fig. S12C). We noted the following changes in these distributions relative to Fig. S12B: (a) the median predictive power for all random gene regressions decreased substantially (i.e., $\bar{R}^2_{CV}$ decreased), indicating that indeed a substantial part of the predictive power of random genes arises due to large correlations with denitrification genes, and (b) denitrification genes now outperformed the typical set of random genes as predictors for even the $r_I$ regression, demonstrating the superiority of denitrification genes as predictors for regressions on all consumer-resource parameters.

#### 2.5 Investigating 16S copy number and genome size as predictors for consumer-resource parameters

We asked whether either 16S rRNA copy number or genome size could serve as better predictor variables for the consumer-resource parameters $\{r_A, \gamma_A, r_I, \gamma_I\}$ than denitrification genes. Previous work demonstrates a positive relationship between 16S copy number and growth rate across diverse taxa in nutrient-replete conditions (Roller et al., 2016; Li et al., 2019), likely because increased ribosome production allows a higher rate of protein synthesis, thereby increasing cell growth rate (Scott et al., 2010). A negative relationship between growth rate and genome size has also been observed (Li et al., 2019). possibly due to a reduced nutrient burden required by smaller genomes (Hessen et al., 2010).

We estimated 16S copy number for all $62$ strains in our library using the 16Stimator pipeline (Perisin et al., 2016) (Table S8). This approach uses Illumina sequencing reads and annotated *de novo* assemblies to compute the coverage ratio of the 16S gene relative to a curated set of single-copy genes. We estimated the genome size for all strains in our library by summing the lengths of all assembled contigs for each strain (Table S8). For our reference strain *Paracoccus denitrificans* ATCC 19367, for which a complete genome assembly shows $3$ copies of the 16S gene and a genome size of $5.24$ mb (Si et al., 2019), we obtain the estimates of $3.36$ 16S copies and a genome size of $5.15$ mb.

We first considered the relationships between 16S copy number, genome size, and the phenotypic parameters $\{r_A, \gamma_A, r_I, \gamma_I\}$ (Fig. S13A and B). We observed a modest but significant (permutation test) positive correlation between 16S copy number and the phenotypic parameter $r_A$ ($\rho = 0.29, p = 0.01$), and significant positive correlations between genome size and $\gamma_A$ ($\rho = 0.50$, $p < 10^{-4}$) and $\gamma_I$ ($\rho = 0.65$, $p < 10^{-4}$). These data suggest that genome size may be a good predictor for the yields $\gamma_A$ and $\gamma_I$.

In order to compare our measurements to previous findings relating growth rate to 16S copy number and genome size, we computed growth rates on nitrate and nitrite as $\mu_* = r_* \gamma_*$ (Fig. S13C). We did not observe a significant positive correlation between either of the growth rates and 16S copy number. This differs from what has been previously observed under aerobic, nutrient replete conditions (Roller et al., 2016; Li et al., 2019). We speculate that this is because growth rates are far from optimal in denitrifying conditions, and the relative benefit of high gene copy number of rRNA may be small. However we observed significant (permutation test) positive correlations between genome size and $\mu_A$ ($\rho = 0.23$, $p = 0.04$), and $\mu_I$ ($\rho = 0.54$, $p < 10^{-4}$). This finding also differs the previous observation of a negative relationship between genome size and growth rate (Li et al., 2019).

We evaluated the potential of 16S copy number and genome size as predictor variables by performing OLS regressions of these predictors on the phenotypic parameters $\{r_A, \gamma_A, r_I, \gamma_I\}$ (Fig. S14A). Regressions on the reduction rate parameters result in poor fits ($R^2 = 0.08$ and $0.06$ for $r_A$ and $r_I$, respectively). Regressions on the yield parameters result in relatively better fits ($R^2 = 0.35$ and $0.44$ for $\gamma_A$ and $\gamma_I$, respectively), driven primarily by the positive correlation between genome size and yield (Fig. S13B). We concluded from this analysis that (a) neither 16S copy number nor genome size are good predictors for the reduction rate parameters, and (b) genome size may be a good predictor for yields $\gamma_A$ and $\gamma_I$, though it is still unclear if genome size holds greater predictive power than the denitrification genes.

We therefore performed a head-to-head comparison between denitrification genes, 16S copy number, and genome size by using all three sets of predictors simultaneously in LASSO regressions on the phenotypic parameters $\{r_A, \gamma_A, r_I, \gamma_I\}$ (4-fold cross-validation, iterated $10^3$ times). We note that, as before, all predictors were standardized to have zero mean and unit variance before fitting. The results of these regressions are shown in Fig. S14B. We observed that: (a) neither 16S copy number nor genome size are assigned large coefficients in any regression, (b) the statistics of fitting and generalization quality ($R^2_{fit}$ and $\bar{R}^2_{CV}$) are essentially the same as those obtained in the original regressions (Fig. 2C to F), and (c) the coefficients for denitrification genes are very similar as those obtained in the original regressions (Fig. 2G to J). We concluded that the denitrification genes hold greater predictive power than 16S copy number and genome size, since the latter predictors are not selected as important variables by LASSO regression.

#### 2.6 Using the consumer-resource model to predict community metabolic dynamics

We extended the consumer-resource model (3) to generate predictions for community (i.e., multi-strain) metabolite dynamics. For an $N$-strain community, the extended model is as follows:

$$\begin{aligned}
\frac{dx_i}{dt} &= \left(\gamma_A r_A^i \frac{A}{K_A + A} + \gamma_I^i r_I^i \frac{I}{K_I + I}\right) x_i \quad \text{for } i = 1, \ldots, N \\
\frac{dA}{dt} &= -\sum_{i=1}^{N} r_A^i \frac{A}{K_A + A} x_i \\
\frac{dI}{dt} &= \sum_{i=1}^{N} \left(r_A^i \frac{A}{K_A + A} - r_I^i \frac{I}{K_I + I}\right) x_i.
\end{aligned} \tag{7}$$

For each strain $i$, which has cell density $x_i$, we have measured the parameters $\{r_A^i, r_I^i, \gamma_A^i, \gamma_I^i\}$. Note that, as before, we fix $K_A = K_I = 0.01\,\mathrm{mM}$ for all strains. This "additive" model sums the independent rate contributions of each strain to the nitrate and nitrite differential equations, in effect assuming that strains only interact via cross-feeding and competition for extracellular nitrate and nitrite. The model *does not* assume that strains interact through Lotka-Volterra-type (quadratic) terms in the cell density equations, nor does it assume that the parameters $\{r_A^i, r_I^i, \gamma_A^i, \gamma_I^i\}$, measured in monoculture, are modulated by the presence of any other strain. Thus equation (7) represents a null model for community metabolite dynamics where each strain in a community behaves as it does in monoculture, and therefore provides a prediction requiring no additional free parameters.

We assembled and cultured simple communities to test the predictions of equation (7), drawing from a representative 12-strain subset of our 62-strain library (Table S9). Specifically, we measured nitrate/nitrite dynamics and endpoint optical density for (a) all 66 2-strain combinations, (b) a random set of 81 3-strain combinations, and (c) controls of each strain in monoculture. All communities/controls were cultured in each of two media conditions, $2\,\mathrm{mM}$ nitrate and $2\,\mathrm{mM}$ nitrite, with 2–3 experimental replicates per condition. Initial cell densities were $\mathrm{OD600}_0 = 0.01$ for each strain in a given community.

We first compared observed endpoint cell densities with the values predicted by the additive model (equation (7)). Since our optical density measurement cannot discern between strains in a mixed population, we computed a model prediction for total endpoint cell density by summing the endpoint cell densities of each strain. We found that endpoint cell densities are well-predicted by equation (7) (Fig. S15). This gives one indication that the additive consumer-resource model in equation (7) accurately predicts community behavior using only information from monocultures.

We next compared the community nitrate and nitrite dynamics measurements with predictions of equation (7). We computed a normalized root-mean-square error (NRMSE) for a given $N$-strain community as

$$NRMSE_{1,\ldots,N} = \frac{RMSE_{1,\ldots,N}}{\left(\frac{1}{N}\sum_{i=1}^{N} RMSE_i^2\right)^{1/2}}, \tag{8}$$

where $RMSE_{1,\ldots,N}$ is the root-mean-square error between measurements and predictions for the community, and $RMSE_i$ is the error for each constituent strain in monoculture, with errors averaged over experimental replicates. We normalized by the monoculture RMSEs of each strain in order to correct for variations in monoculture fit quality. Values of NRMSE less than or equal to $1$ indicate high quality predictions of community metabolite dynamics, with prediction errors similar to those of the constituent monocultures, while values of NRMSE much greater than $1$ indicate low quality predictions.

The NRMSEs for all pairs of $12$ strains cultured in separate nitrate and nitrite medium conditions are shown in Fig. S16A and B. For most pair communities, we inferred from typically low values of NRMSE that equation (7) made high quality predictions in most cases. The apparent exception to this occurs for "Nar + Nir" communities (comprising any combination of Nar strain and Nir strain) initialized with nitrate (Fig. S17), for which NRMSE values appeared much higher than for any other group. We performed permutation tests on the mean NRMSE values within phenotypic groups to determine whether, in fact, the "Nar + Nir" group has significantly higher NRMSE than other groups ($10^5$ permutations). Since we performed $10$ hypothesis tests in this manner, a Bonferroni-corrected threshold of $p = 0.05/10 = 0.005$ was used for testing at the $5\%$ significance level. We find that indeed only the "Nar + Nir" group in nitrate medium conditions has a significantly large mean NRMSE value ($p < 10^{-5}$).

The NRMSEs for $81$ randomly-selected 3-strain communities cultured in separate nitrate and nitrite medium conditions are shown in Fig. S18. As with pair communities, we observed that most 3-strain communities have low values of NRMSE indicating that equation (7) made high quality predictions of nitrate and nitrite dynamics. We observed also that most apparent failures of the model predictions occur when a 3-strain community contains both a Nar strain and a Nir strain in the nitrate medium condition.

#### 2.7 Correcting for interactions between Nar and Nir strains

The additive $N$-strain consumer-resource model (7) generally failed to predict the dynamics of communities initialized with nitrate that included both a Nar and a Nir strain (see plots of all Nar + Nir pairs in Fig. S17). In every instance, the nitrate reduction rate of the Nar strain appears to be diminished relative to the rate in monoculture, and in some instances the nitrite reduction rate for the Nir strain appears to be increased relative to monoculture. Since Nar and Nir strains utilize different electron acceptors and therefore do not plausibly compete for any resource under these experimental conditions, we concluded that interactions aside from resource competition must be taking place.

We therefore asked whether the predictions of the additive model in equation (7) can be improved by correcting for interactions between Nar and Nir strains. To do this, we refit the nitrate and nitrite reduction parameters $r_A^i$ and $r_I^j$ of the Nar strain $i$ and the Nir strain $j$, respectively, using the measured metabolite dynamics data from the pair cultures of these strains. We focused on identifying changes to the reduction rate parameters rather than the yields ($\gamma_A^i$ and $\gamma_I^j$) because the additive model accurately predicted endpoint optical densities in all pair communities (Fig. S15C and D), and therefore there was no evidence that interactions between Nar and Nir strains involved a change in yields.

To refit nitrate and nitrite reduction rate parameters for each Nar + Nir pair culture, we:

1. Refit $r_A^i$ for the Nar strain $i$, holding all other parameters fixed. We then computed the NRMSE for the resulting prediction.

2. Refit both $r_A^i$ for the Nar strain $i$ and $r_I^j$ for the Nir strain $j$, holding all other parameters fixed. We again computed the NRMSE for the resulting prediction.

3. If the NRMSE obtained in step 2 was more than $10\%$ smaller than the NRMSE in step 1, then we accepted the refit parameters obtained in step 2, denoting these values as $\tilde{r}_A^{i,j}$ and $\tilde{r}_I^{j,i}$. Otherwise, we accepted the refit parameter obtained in step 1 as $\tilde{r}_A^{i,j}$, and let $\tilde{r}_I^{j,i} = r_I^j$.

Refitting was performed as described above for equation (3). In step 3, we compared the improvements in fit quality obtained by refitting only $r_A$ versus refitting both $r_A$ and $r_I$ in order to identify cases where increases in nitrite reduction rate were necessary to improve fits of equation (7) to the data.

The results of this refitting procedure are shown in Fig. S19. These results demonstrate that, in all cases, the nitrate reduction rate of the Nar strain was slowed ($\tilde{r}_A < r_A$) in Nar + Nir pair culture. Additionally, in several cases the nitrite reduction rate of the Nir strain was increased ($\tilde{r}_I > r_I$) in Nar + Nir pair culture. We note that each Nar strain is affected in essentially the same way by every Nir strain (e.g., the nitrate reduction rate of Nar strain PAR01 is reduced by approximately $50\%$ in every pair culture with a Nir strain).

We then used these refit parameter values in equation (7) to generate new predictions for 3-strain communities containing at least one Nar + Nir pair. To do so, we employed the following rules for replacing $r_A$ and $r_I$ in equation (7), which depend on the number of Nar and Nir strains in the community:

1. One Nar strain $i$ and one Nir strain $j$: $r_A^i = \tilde{r}_A^{i,j}$, $r_I^j = \tilde{r}_I^{j,i}$.

2. Two Nar strains $i$ and $j$ and one Nir strain $k$: $r_A^i = \tilde{r}_A^{i,k}$, $r_A^j = \tilde{r}_A^{j,k}$, $r_I^k = \text{mean}(\tilde{r}_I^{k,i}, \tilde{r}_I^{k,j})$.

3. One Nar strain $i$ and two Nir strains $j$ and $k$: $r_A^i = \text{mean}(\tilde{r}_A^{i,j}, \tilde{r}_A^{i,k})$, $r_I^j = \tilde{r}_I^{j,i}$, $r_I^k = \tilde{r}_I^{k,i}$.

The results of using refit Nar and Nir parameters to predict 3-strain cultures are shown in Fig. 4D (yellow points). We observed that correcting predictions using refit $r_A$ and $r_I$ parameters substantially improves prediction quality, reducing median NRMSE values from $2.1$ to $0.79$. These results do not appreciably change if a min or max function is used instead of mean in rules 2 and/or 3.

#### 3 Supplemental Figures

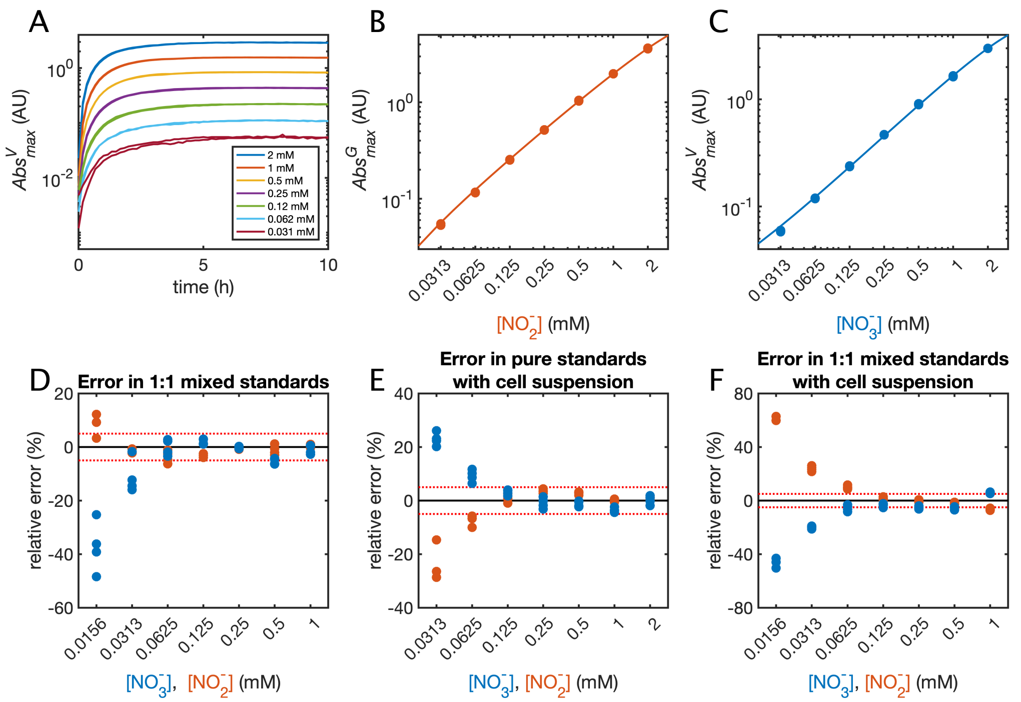

Figure S1: **Nitrate/nitrite assay quality control.** (**A**) Nitrate standards (in duplicate) were incubated with Griess reagents and $VCl_3$ solution at 30 °C, and the blank-corrected maximum absorbances ($Abs^V_{max}$) were recorded and plotted over time. (**B** and **C**) Example nitrite and nitrate standard curves. Measured absorbances (points) of nitrite and nitrate standards were used to fit equations (1) and (2), resulting in the plotted curves. Absorbance values are given in dimensionless absorbance units (AU). (**D** and **F**) Standard curves were used to infer concentrations of nitrate and nitrite in additional standards (four replicates of each condition). The relative errors of nitrate and nitrite measurements are shown in blue and orange points, respectively. The dashed red lines indicate ± 5% relative error. Panel **D** shows relative errors for nitrate and nitrite standards mixed in a 1:1 ratio. Panel **E** shows relative errors for separate nitrate and nitrite standards that have been mixed with a cell suspension. Panel **F** shows relative errors in 1:1 mixtures of nitrate and nitrite that have been mixed with a cell suspension.

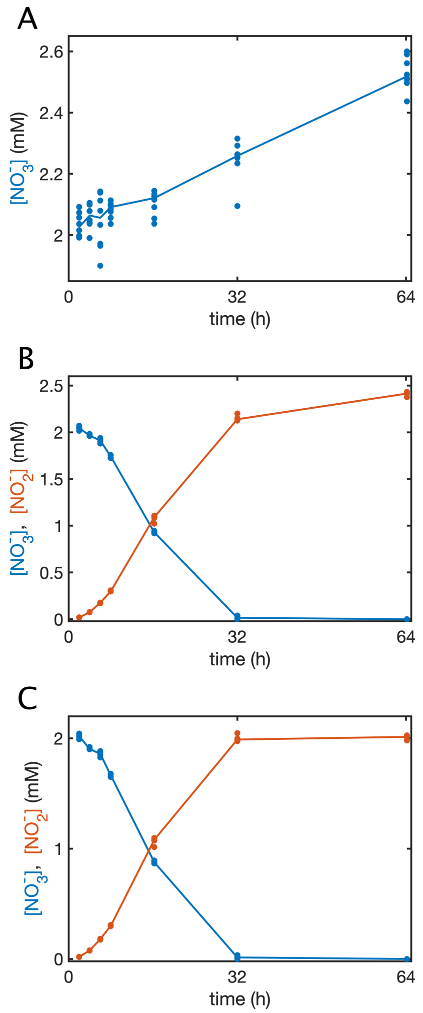

Figure S2: **Correcting nitrate/nitrite dynamics data for evaporation effects.** (**A**) Nitrate dynamics data for $8$ uninoculated blanks initialized with $2\,\text{mM}$ $\text{NO}_3^-$. (**B** and **C**) Example nitrate and nitrite dynamics data for the Nar strain *Variovorax* sp. VRV01 ($N = 4$ replicates). Panel **B** shows raw data and panel **C** shows evaporation-corrected data. Points show measured concentrations and lines connect median values at each time point.

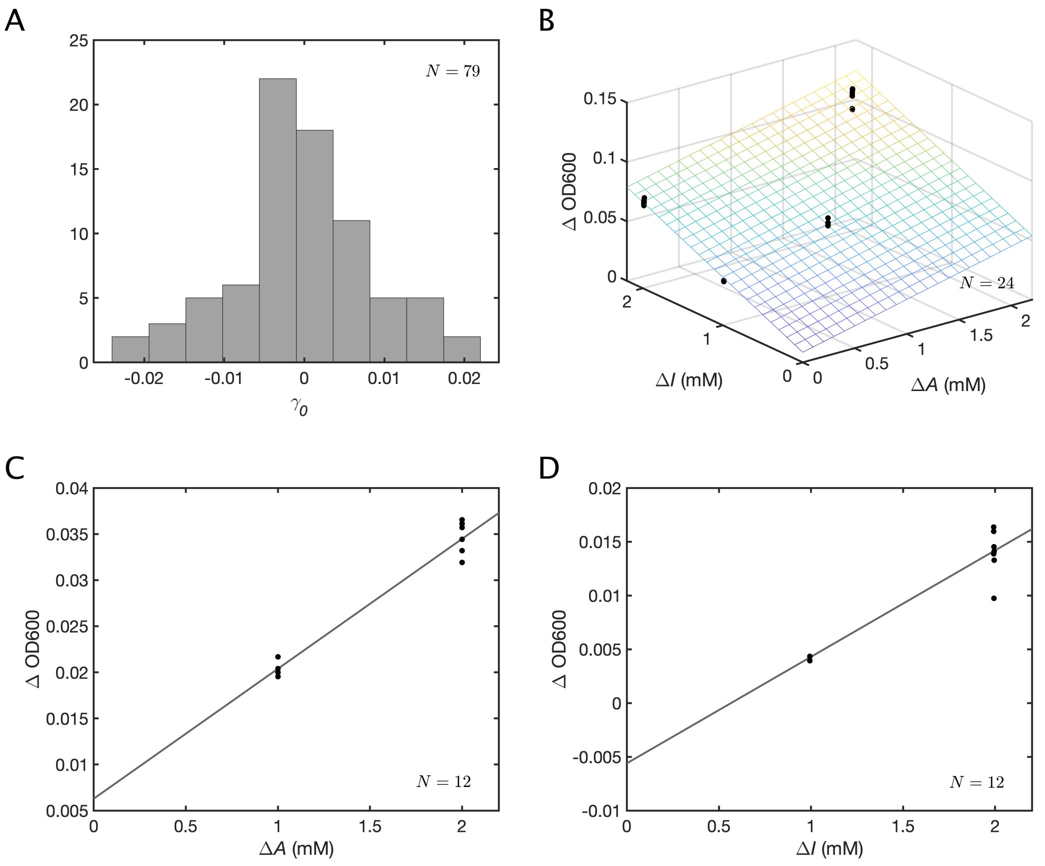

Figure S3: **Fitting consumer-resource yield parameters.** (**A**) Histogram showing the distribution of yield intercept ($\gamma_0$) values for $79$ strains ($78$ isolates and one reference strain *Paracoccus denitrificans* ATCC 19367). Units are dimensionless absorbances at $600\,\mathrm{nm}$, path length normalized to $1\,\mathrm{cm}$. (**B** to **D**) Examples of yield parameter fits using data obtained in different growth conditions. Points show observed values of $\Delta$OD600 from different conditions where different amounts of nitrate and nitrite are reduced ($\Delta A$ and $\Delta I$, respectively), with $4$ replicates used in each condition. The plane/lines show the least-squares fits of the data to equation (4). Panel **B** shows the yield fit for the Nar/Nir strain *P. denitrificans* ATCC 19367. Panel **C** shows the yield fit for the Nar strain *Raoultella* sp. RLT01. Panel **D** shows the yield fit for the Nir strain *Pseudomonas* sp. PDM13.

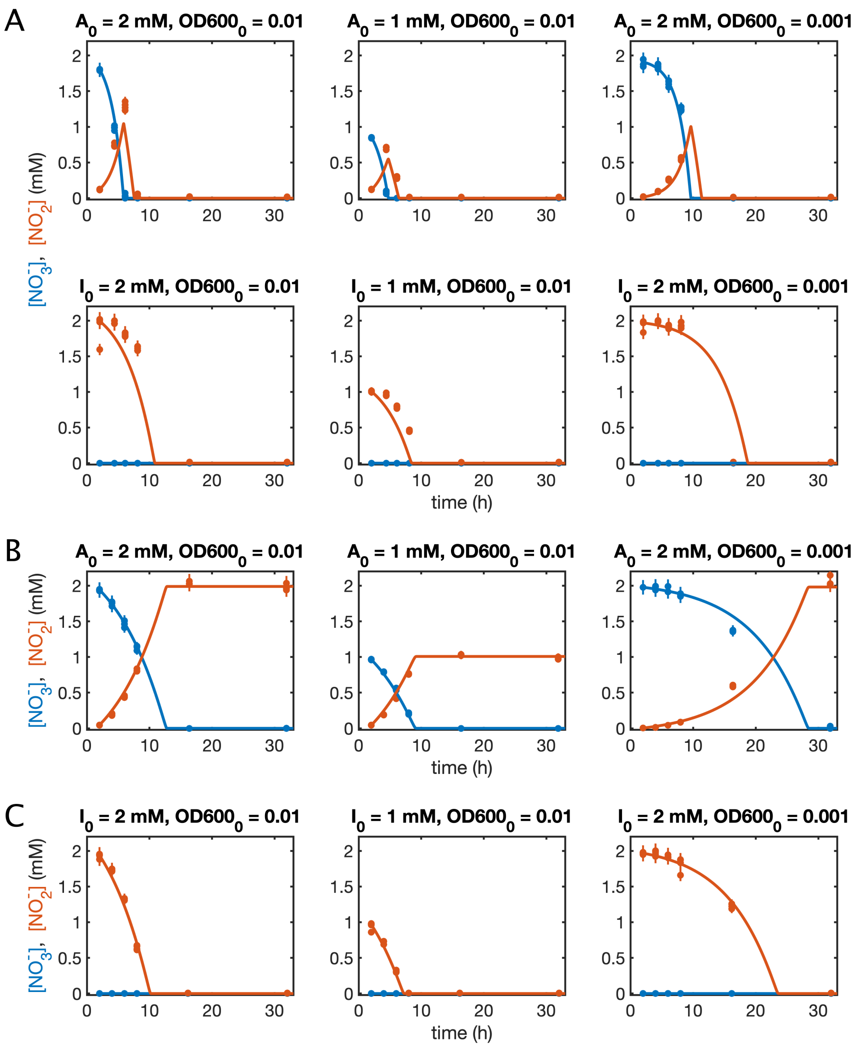

Figure S4: **Fitting consumer-resource reduction rate parameters to nitrate/nitrite dynamics data.** (**A** to **C**) Example global fits of the consumer-resource model (3) to nitrate and nitrite dynamics data from different experimental conditions. Points show measured concentrations of nitrate and nitrite, and curves show optimal model fits. Panel **A** shows global fits for the Nar/Nir strain *P. denitrificans* ATCC 19367 using data collected from six different experimental conditions. Panel **B** shows global fits for the Nar strain *Raoultella* sp. RLT01 using data from three different experimental conditions. Panel **C** shows global fits for the Nir strain *Pseudomonas* sp. PDM13 using data from three different experimental conditions. Four replicates were used in each experimental condition.

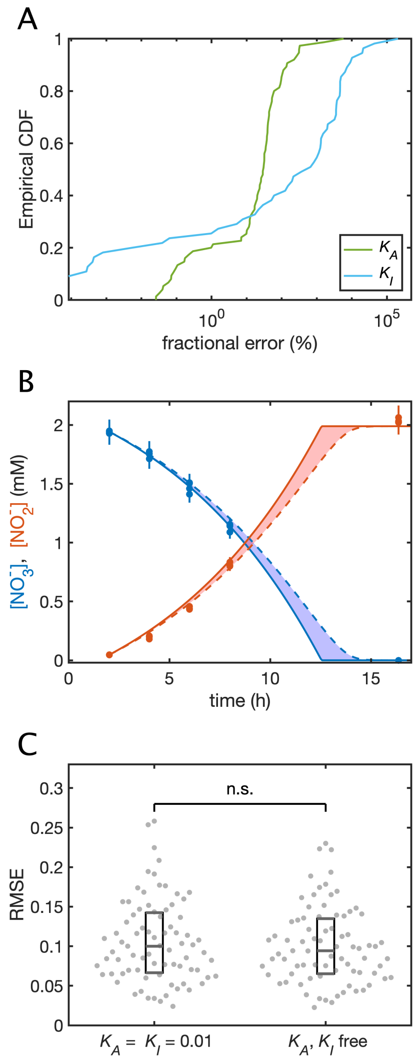

Figure S5: **Justifying the choice of fixed values of the consumer-resource substrate affinity parameters.** (**A**) Distributions of fractional errors (%) for the affinity parameters $K_A$ and $K_I$, computed via nonparametric bootstrap with $K_A, K_I$ constrained during fitting between $0.001$ and $10$. Fractional error is defined as the ratio of the interquartile range obtained via bootstrapping to the value of the parameter obtained using a standard fit to all experimental data. (**B**) Example nitrate and nitrite dynamics for the Nar strain *Raoultella* sp. RLT01. Solid lines show the fit to equation (3) holding $K_A$ fixed at $0.001$, while dashed lines show fits holding $K_A = 0.1$. Points show measured concentrations of nitrate and nitrite. (**C**) Comparison of model fit errors (RMSE) for $N = 79$ denitrifying strains. Fits that hold $K_A = K_I = 0.01$ and fits that take $K_A$ and $K_I$ as free fitting parameters are compared. A two-sample Kolomogorov-Smirnov test accepts the null hypothesis that underlying distributions for the two samples are the same ($p = 0.97$). Box plots indicating quartiles of each distribution are shown.

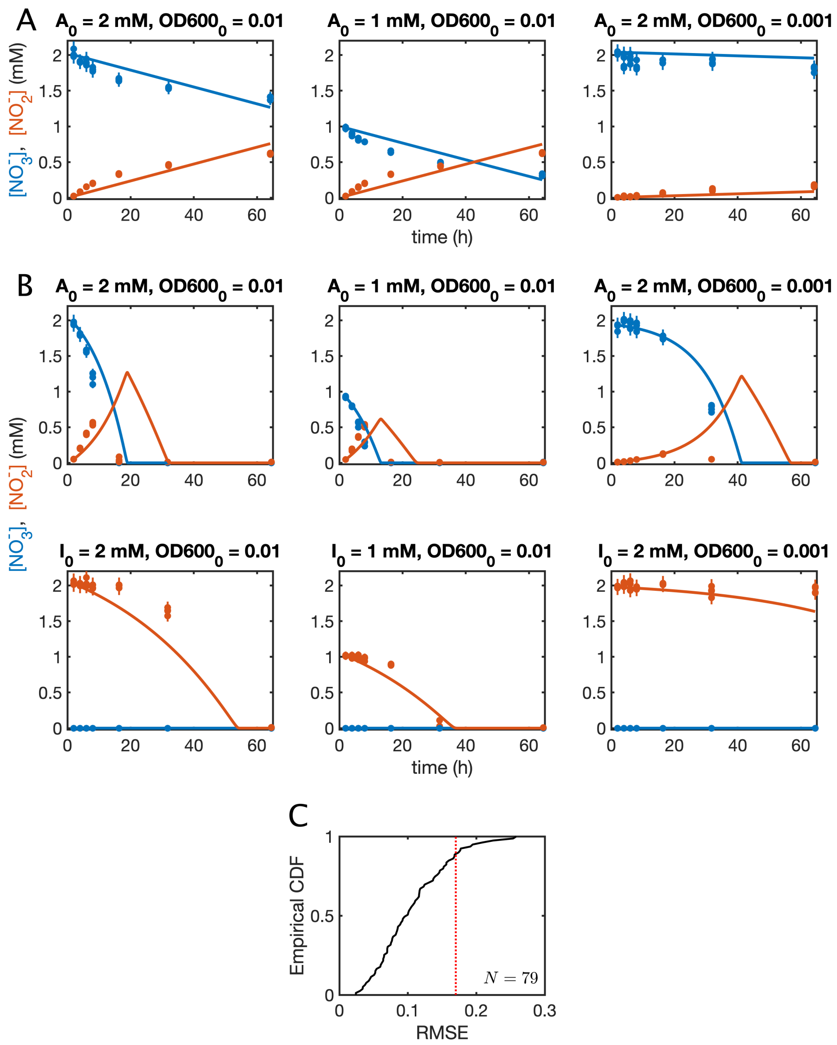

Figure S6: **Excluding poorly-fitting strains from the isolate library.** (**A**) Example Nar strain for which nitrate concentrations appear to asymptotically approach a nonzero value. Points show measured concentrations of nitrate and nitrite, and curves show optimal model fits to three experimental conditions. (**B**) Example Nar/Nir strain with a low quality model fit (RMSE $= 0.173$). Points show measured concentrations of nitrate and nitrite, and curves show optimal model fits to six experimental conditions. (**C**) Empirical CDF of RMSE values with a threshold drawn at $0.17$, above which strains are excluded from further analysis. RMSEs were evaluated between observed data and model solutions across all experimental conditions.

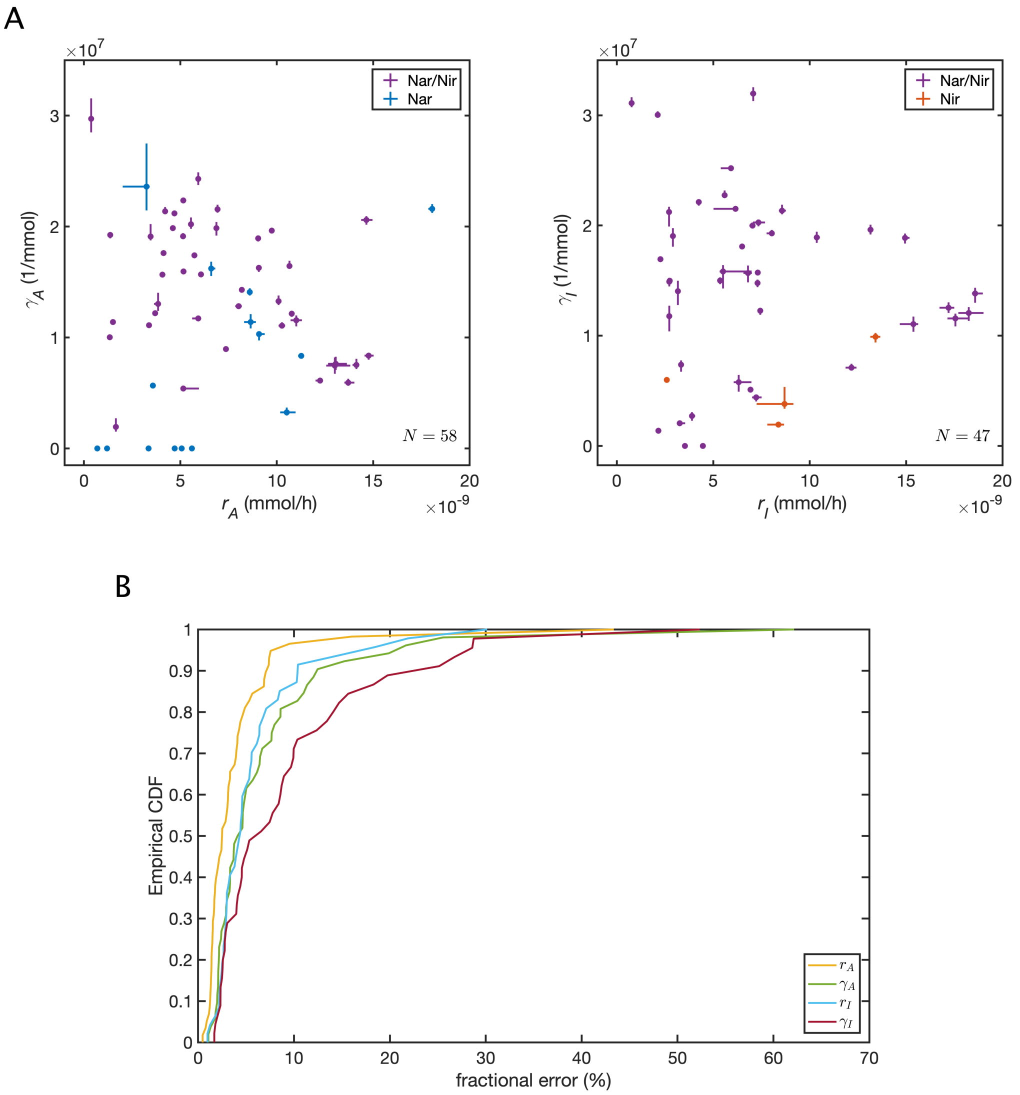

Figure S7: **Covariation and errors for the consumer-resource parameters.** (**A**) Observed variation in measured metabolite dynamics parameters for $62$ denitrifying strains ($58$ nitrate reducers and $47$ nitrite reducers). Values for Nar/Nir, Nar, and Nir strains are shown in purple, blue, and orange respectively. Error bars show the 25th to 75th percentile estimates computed via nonparametric bootstrap. (**B**) Distributions of fractional errors (%) for each consumer-resource parameter. Fractional error is defined as the ratio of the interquartile range obtained via bootstrapping to the value of the parameter obtained using a standard fit to all experimental data.

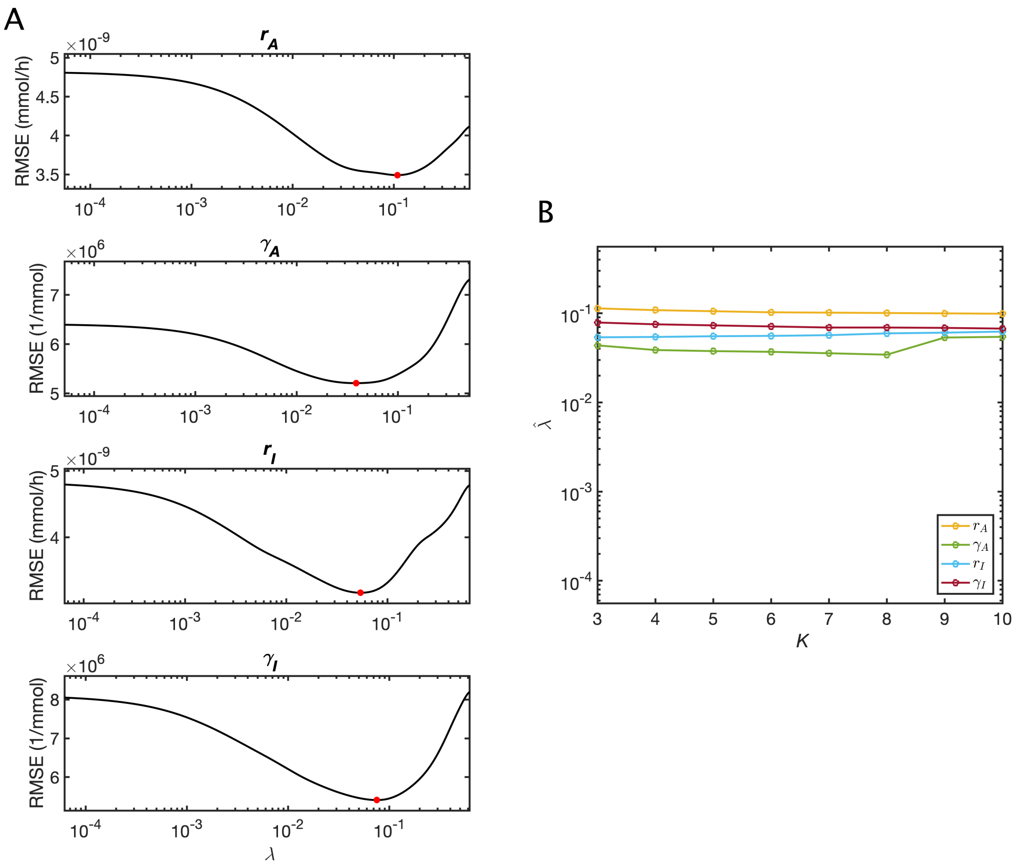

Figure S8: **Using cross-validation for hyperparameter selection.** (**A**) Plots showing the hyperparameter value $\lambda$ versus root-mean-square prediction error using iterated 4-fold cross-validation ($M = 10^4$ iterations) for regressions on each of the consumer-resource parameters. Minimum RMSE values that determine $\lambda = \hat{\lambda}$ are indicated in red points. (**B**) Values of $\hat{\lambda}$ selected via iterated $K$-fold cross-validation ($M = 10^4$ iterations) for different values of $K$ for each of the consumer-resource parameters.

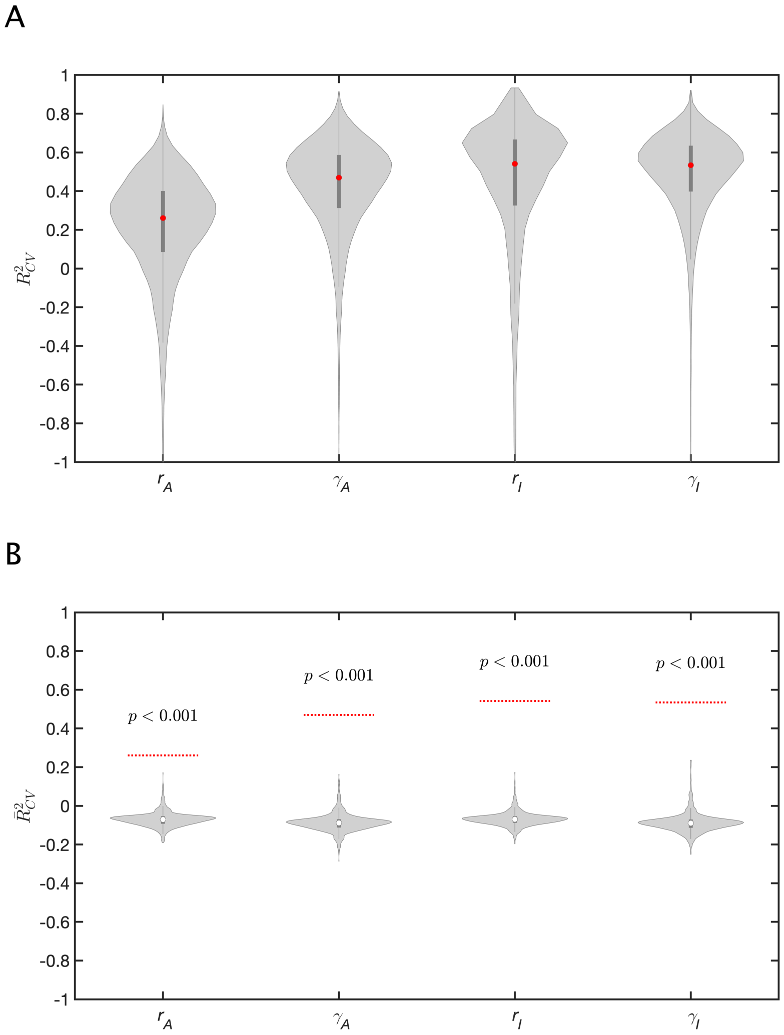

Figure S9: **Estimating out-of-sample performance.** (**A**) Violin plots showing the distributions of coefficients of determination obtained during iterated cross-validation ($R^2_{CV}$) for LASSO regressions on the consumer-resource parameters $\{r_A, \gamma_A, r_I, \gamma_I\}$. Values were obtained at the optimal hyperparameter value determined by cross-validation, $\lambda = \hat{\lambda}$. Median values of these distributions, denoted $\bar{R}^2_{CV}$ are indicated in red. (**B**) Violin plots showing the null distributions of $\bar{R}^2_{CV}$ obtained via a permutation test ($10^3$ permutations), under the null hypothesis that there is no relationship between predictor and response. Dashed red lines indicate the observed values of $\bar{R}^2_{CV}$ on the unpermuted data (the red points in panel **A**). In all cases, $p$-values are $< 0.001$.

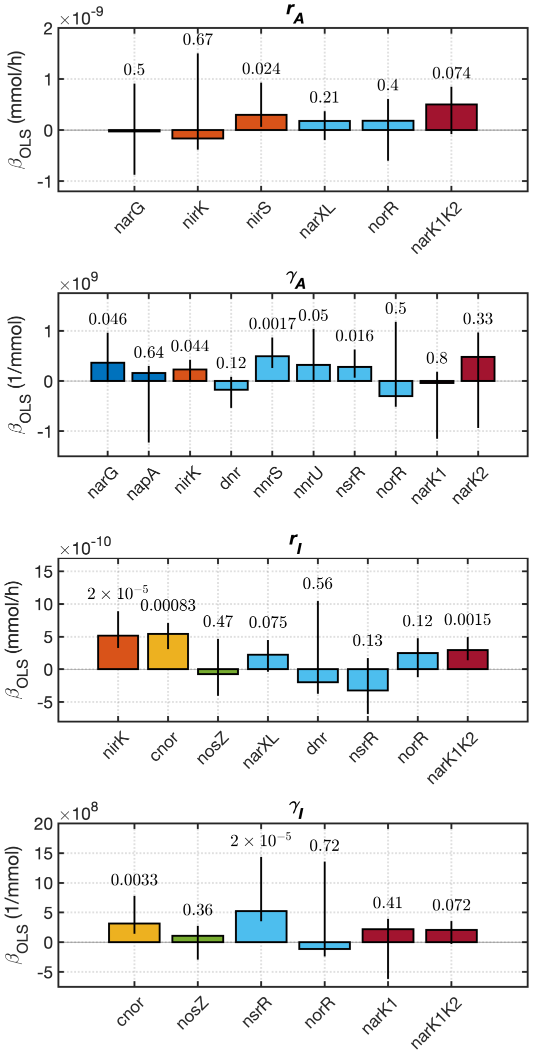

Figure S10: **Post-selection inference on the nonzero coefficients determined by LASSO.** OLS regression coefficients on consumer-resource parameters $\{r_A, \gamma_A, r_I, \gamma_I\}$ using only variables selected by LASSO regressions, along with $90\%$ confidence intervals and $p$-values computed using the selectiveInference package for R.

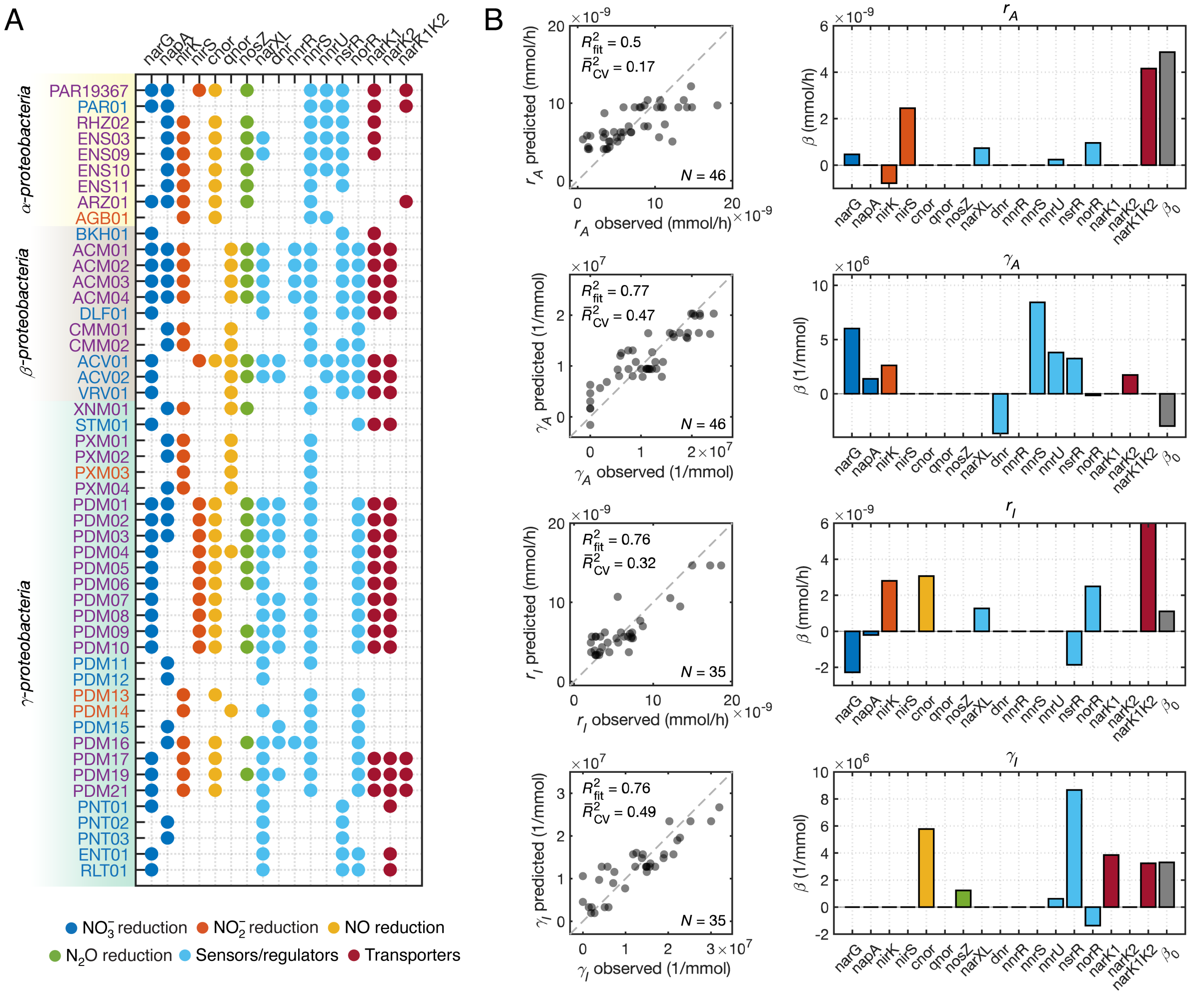

Figure S11: **Quantifying the impact of phylogenetic correlations on denitrification gene regressions.** (**A**) Denitrification gene presence/absence for a subsampled library of 50 denitrifying strains. Clades containing strains with identical 16S rRNA sequences were "collapsed" onto one randomly-selected representative, resulting the removal of 12 strains from the full 62-strain library. (**B**) Regression applied to each consumer-resource parameter using the 50-strain subsampled dataset resulted in predicted values of the parameters for each strain, which are plotted against observed values, and fitted values of the vector $\vec{\beta}$, whose coefficients, along with the intercept $\beta_0$, are shown in the bar charts. The in-sample coefficients of determination for these data ($R^2_{\text{fit}}$) and the out-of-sample coefficients of determination estimated via iterated cross-validation ($\bar{R}^2_{\text{CV}}$) are shown.

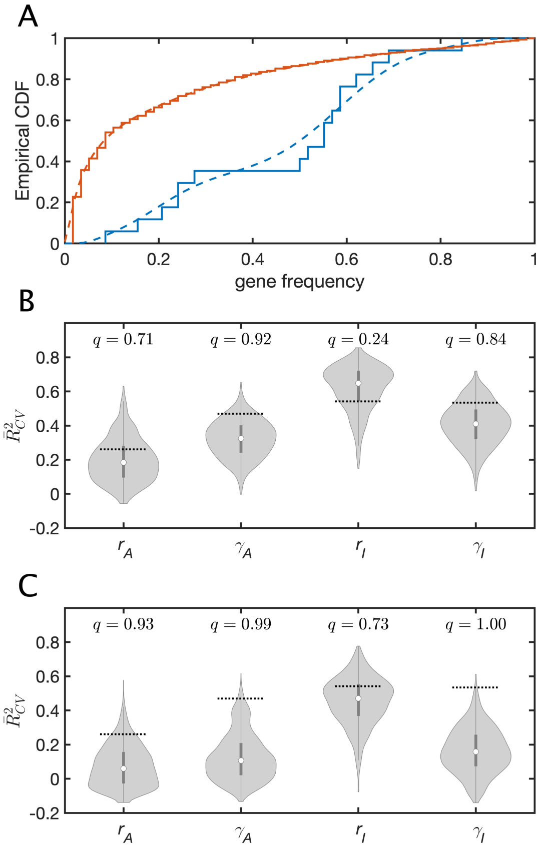

Figure S12: **Randomly-selected genes as alternative predictors for consumer-resource parameters.** (**A**) Distributions of gene presence frequency (fraction of strains that possess a given gene) for denitrification-related genes (blue) and the distribution of gene presence frequency for all other annotated genes (red), both in the ensemble of the $58$ nitrate-reducing strains. Solid lines show empirical CDFs of gene frequencies, and dashed lines show kernel density estimates of these distributions (bandwidth = $0.4$). (**B** and **C**) Distributions of $\bar{R}^2_{CV}$ values obtained by regressing each of the consumer-resource parameters onto the presence/absence of sets of randomly-selected genes ($10^3$ sets of random genes per consumer-resource parameter). Panel **B** shows results for genes randomly selected from the set of all annotated genes across strains in our library, while panel **C** shows results for genes randomly selected from the set of all annotated genes excluding those that have large and significant correlation ($|\rho| \geq 0.5$) with any denitrification genes. Dashed lines indicate $\bar{R}^2_{CV}$ values obtained in regressions onto the presence/absence of denitrification related genes (the same values are shown in both panels), with the corresponding quantile values ($q$) shown above.

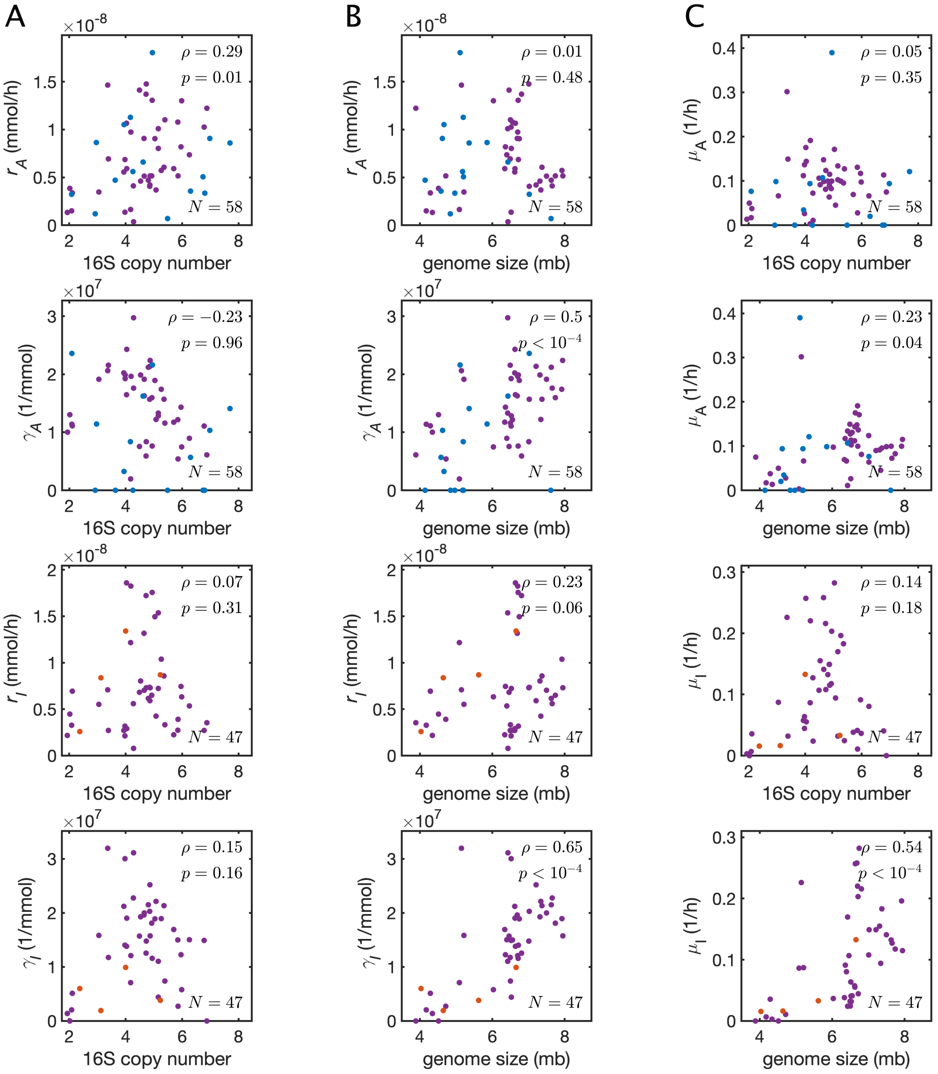

Figure S13: **Correlations between 16S copy number, genome size and the consumer-resource parameters.** (**A** and **B**) Scatter plots of the consumer-resource parameters $\{r_A, \gamma_A, r_I, \gamma_I\}$ and 16S copy number (panel **A**) , and genome size (panel **B**). (**C**) Scatter plots of the growth rates, 16S copy number, and genome size. Growth rates on either nitrate or nitrite are computed as $\mu_* = r_* \gamma_*$. In all plots, values for Nar/Nir, Nar, and Nir strains are shown in purple, blue, and orange respectively. Pearson correlations and $p$-values determined via a permutation test are shown.

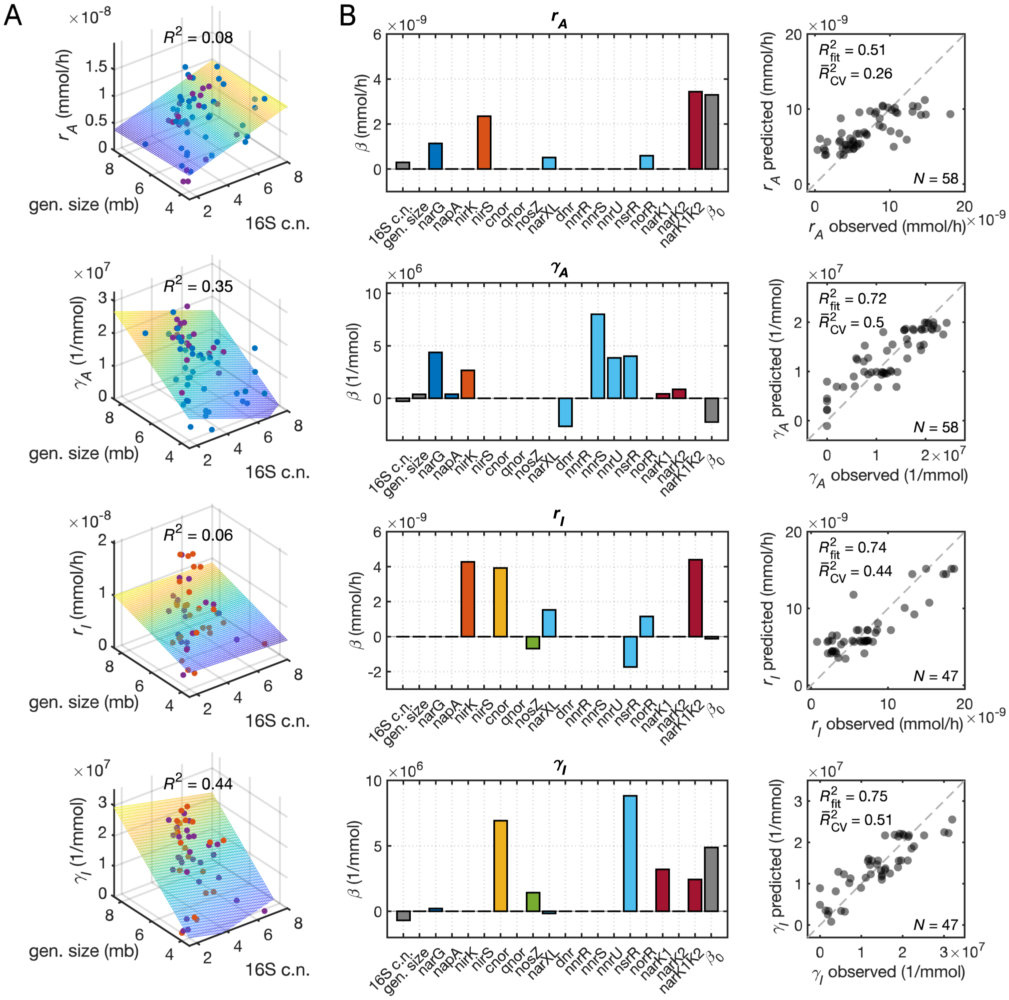

Figure S14: **16S copy number and genome size as alternative predictors for consumer-resource parameters.** (**A**) Results from OLS regressions on consumer-resource parameters $\{r_A, \gamma_A, r_I, \gamma_I\}$ using only 16S rRNA copy number and genome size as predictors. Coefficients of determination ($R^2$) for observed versus predicted values are shown. (**B**) Results from LASSO regressions on phenotypic parameters $\{r_A, \gamma_A, r_I, \gamma_I\}$, where 16S rRNA copy number, genome size, and denitrification gene presence/absence are all simultaneously used as predictors. Regression coefficients $\vec{\beta}$, along with the intercept $\beta_0$, are shown in the bar charts. The scatter plots show observed values of parameters versus values predicted by the regression. The in-sample coefficients of determination for these data ($R^2_{\text{fit}}$) and the median out-of-sample coefficients of determination estimated via iterated cross-validation ($\bar{R}^2_{\text{CV}}$) are shown.

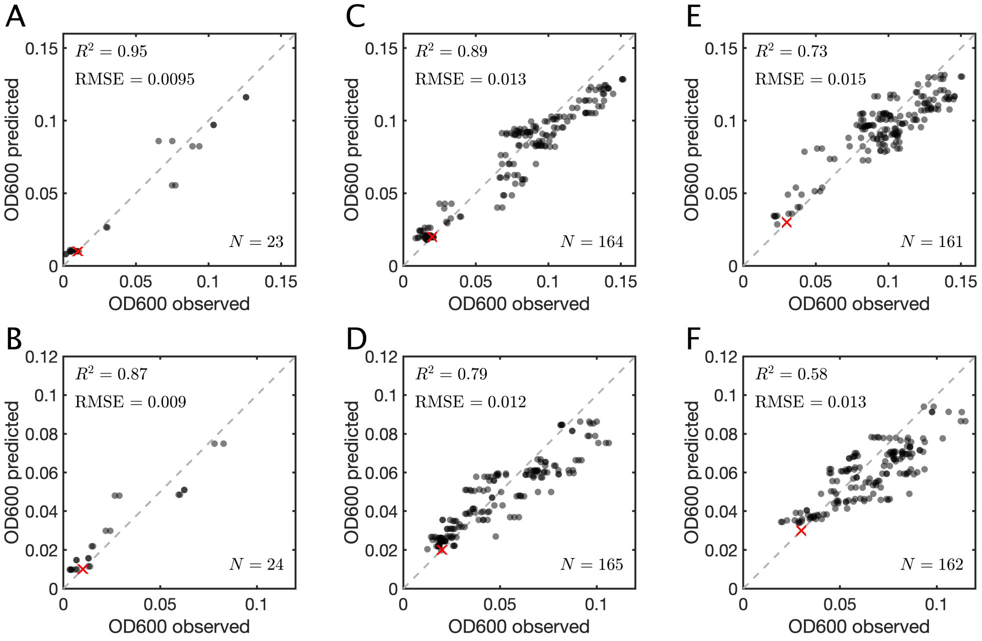

Figure S15: **Accuracy of consumer-resource endpoint cell density predictions.** (**A** to **F**) Comparisons between observed endpoint optical densities and values predicted by the additive consumer-resource model (7), where the latter was obtained by summing the endpoint cell densities of each strain in the community. Comparisons are shown for monoculture controls (**A** and **B**), 2-strain communities (panels **C** and **D**), and 3-strain communities (panels **E** and **F**), in $2\,\text{mM}$ nitrate (panels **A**, **C**, and **E**) and $2\,\text{mM}$ nitrite (panels **B**, **D**, and **F**) media conditions. Initial optical densities are indicated (red cross), and coefficients of determination $R^2$ and root-mean-square errors (RMSE) for observed versus predicted values are shown.

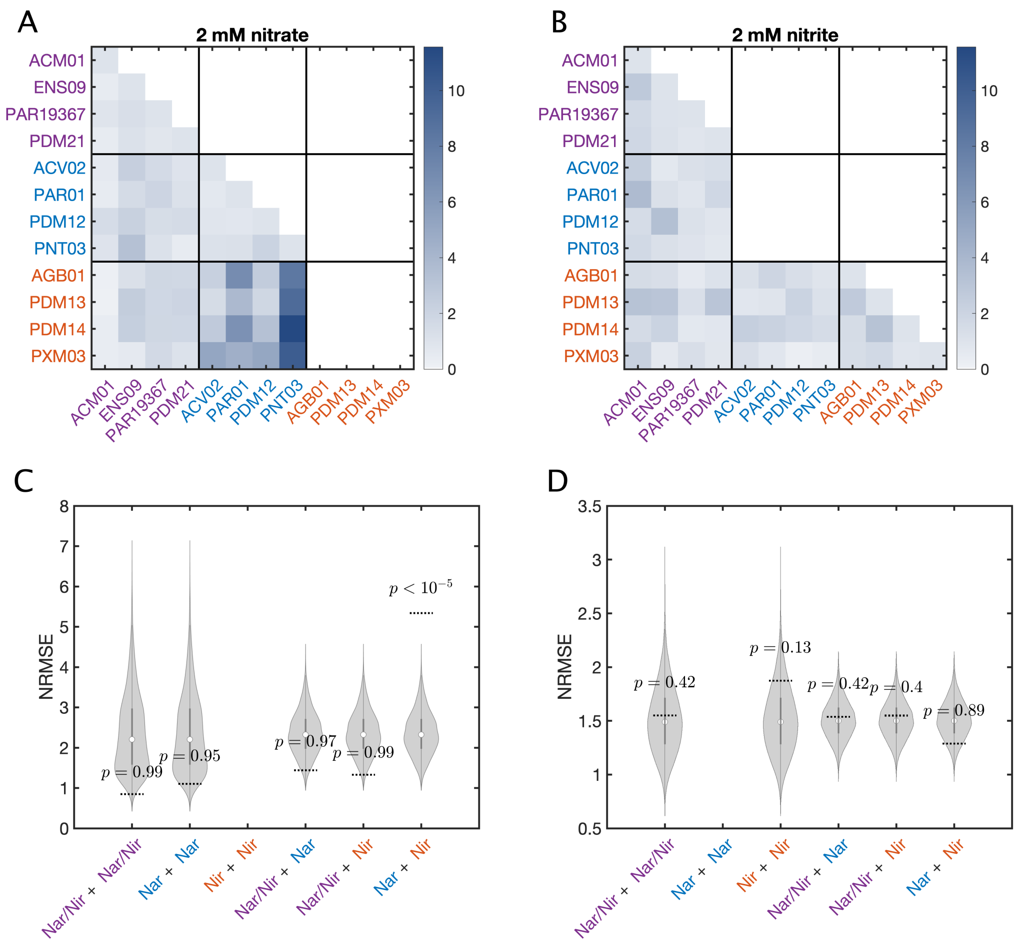

Figure S16: **Accuracy of consumer-resource nitrate/nitrite dynamics predictions for pair communities.** (**A** and **B**) Normalized RMSE (NRMSE) values comparing measurements and predictions for nitrate and nitrite dynamics in all pair communities of 12 strains. Values are grouped according to the constituent phenotypes in a community, with Nar/Nir strains are labelled in purple, Nar strains in blue, and Nir strains in orange. Panel **A** shows NRMSEs for communities initialized with $2\,\mathrm{mM}$ nitrate (with "Nir + Nir" communities omitted, since Nir strains do not utilize nitrate as an electron acceptor). Panel **B** shows NRMSEs for communities initialized with $2\,\mathrm{mM}$ nitrite (with "Nar + Nar" communities omitted, since Nar strains do not utilize nitrite as an electron acceptor). (**C** and **D**) Hypothesis testing on means of NRMSEs grouped by constituent phenotypes (e.g., "Nar/Nir + Nar/Nir" communities, etc.). Null distributions for group means are generated via permutation of group labels for each pair community ($10^5$ permutations), and are compared with the observed mean NRMSEs in each group (dashed lines). A Bonferroni-corrected threshold for 5% significance over 10 hypothesis tests is 0.005. Note that some groups contain only 6 NRMSE values (e.g., "Nar/Nir + Nar/Nir" communities), while others contain 16 NRMSE values (e.g., "Nar/Nir + Nar" communities). Panel **C** shows hypothesis testing on the NRMSE values shown in panel **A** (communities initialized with nitrate), while panel **D** shows inference on the values shown in panel **B** (communities initialized with nitrite).

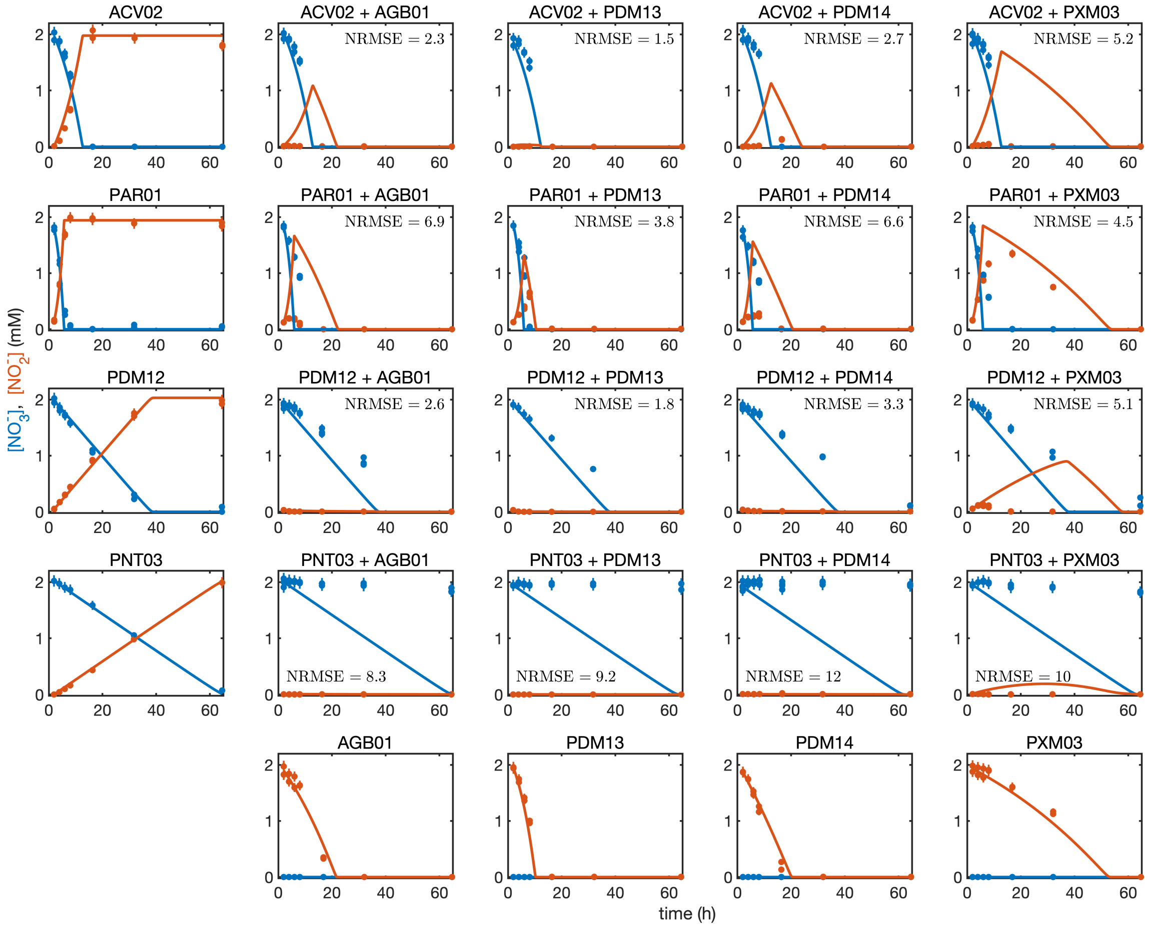

Figure S17: **Nitrate/nitrite dynamics for Nar + Nir pair cultures.** Monoculture controls are shown for Nar strains (left column) and Nir strains (bottom row). Points show measured concentrations of nitrate and nitrite, and curves show predictions of equation (7). NRMSE values for pair cultures are shown. 2–3 experimental replicates are used for each combination of strains.

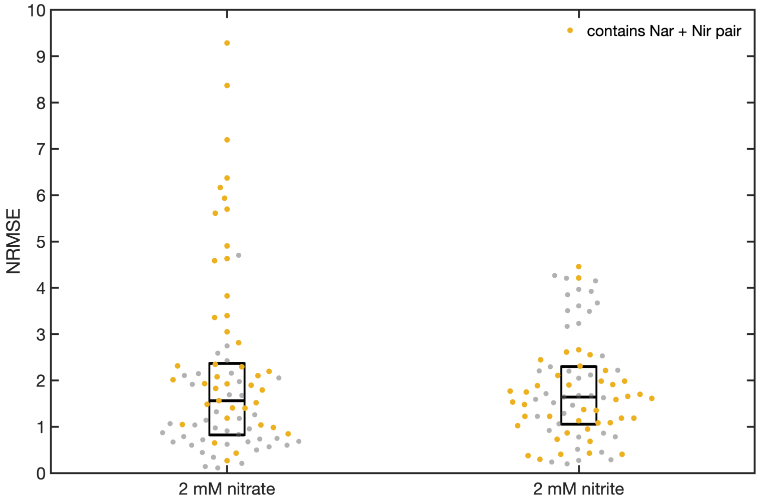

Figure S18: **Accuracy of consumer-resource nitrate/nitrite dynamics predictions for 3-strain communities.** Distributions of normalized RMSE (NRMSE) values comparing measurements and predictions for nitrate and nitrite dynamics in random 3-strain communities ($N = 81$). Communities were separately initialized with $2\,\mathrm{mM}$ nitrate and $2\,\mathrm{mM}$ nitrite. Points in yellow indicate values for communities that contain both a Nar strain and a Nir strain. Box plots indicating quartiles of each distribution are shown.

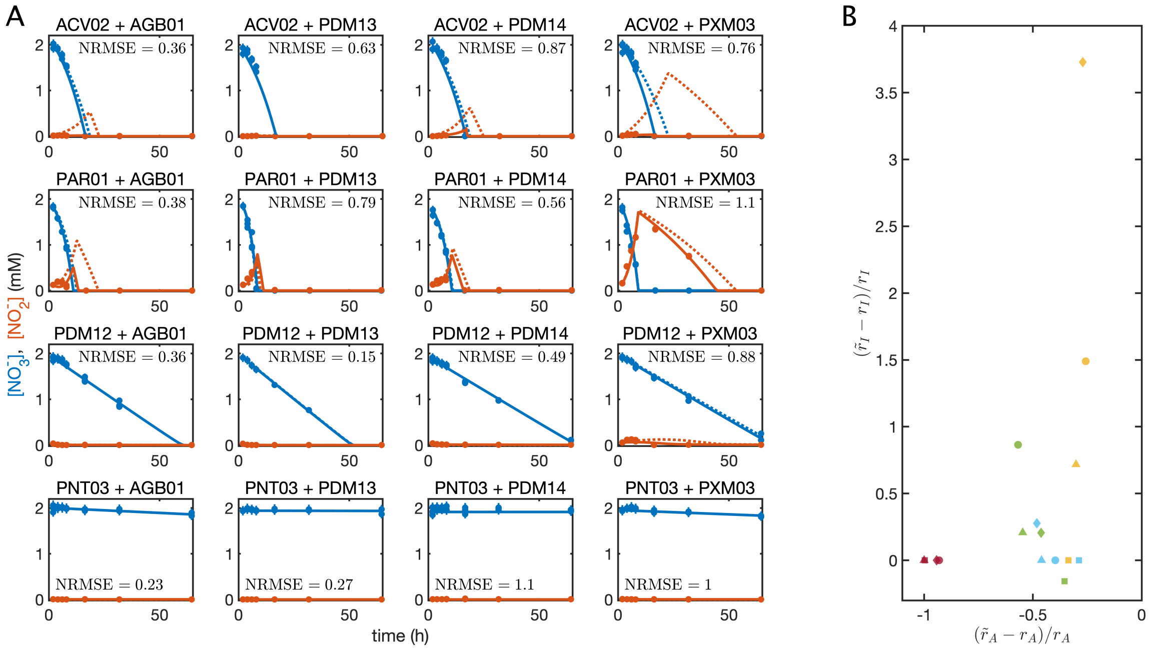

Figure S19: **Refitting model parameters for Nar + Nir pair cultures.** (**A**) The $r_A$ parameter for the Nar strain and/or the $r_I$ parameter for the Nir strain were refit to the Nar + Nir pair culture measurements to obtain new parameter values $\tilde{r}_A$ and $\tilde{r}_I$, respectively. In some cases (panels with only solid lines) it was only necessary to refit the $r_A$ parameter to obtain a good model fit. In other cases (panels with both solid and dashed lines) refitting only $r_A$ produced poor model fits (dashed lines), and so both $r_A$ and $r_I$ were refit (solid lines). NRMSE values for refit models are indicated. 2–3 experimental replicates are used for each combination of strains. (**B**) Relative changes in $r_A$ and $r_I$ parameters. Nar strains are coded by color: ACV02 (yellow), PAR01 (green), PDM12 (blue), and PNT03 (red). Nir strains are coded by symbol: AGB01 (circle), PDM13 (square), PDM14 (triangle), PXM03 (diamond). NRMSE values for pair cultures are shown.

### 4 Supplemental Tables

Table S1: Summary of findings from previous studies comparing the nitrate reduction activity of the membrane-bound nitrate reductase (*NarG*) and the periplasmic nitrate reductase (*NapA*).

| Finding | Ref. |
|---|---|
| Whole cell nitrate reduction activity assay comparing *NarG* and *NapA* knockout mutants of *Escherichia coli* K-12 found that reduction is faster for $\Delta$*NapA* (*NarG*+) mutant than for $\Delta$*NarG* (*NapA*+). | (Stewart et al., 2002) |
| Whole cell nitrate reduction activity assay comparing *NarG* and *NapA* knockout mutants of *Pseudomonas aeruginosa* PAO1 found that reduction is faster for $\Delta$*NapA* (*NarG*+) mutant than for $\Delta$*NarG* (*NapA*+). | (Van Alst et al., 2009) |
| Comparing nitrate reduction activity in soluble (*NapA*) versus membrane (*NarG*) fractions of cell lysate showed higher activity in membrane fraction for both *Pseudomonas* sp. MT1 and *Pseudomonas stutzeri* IFO 14165. | (Ikeda et al., 2009) |
| Comparing nitrate reduction activity in soluble (*NapA*) versus membrane (*NarG*) fractions of cell lysate showed higher activity in membrane fraction for *Thiosphaera pantotropha* LMD 82.5 | (Bell et al., 1990) |
| Comparing nitrate reduction activity in soluble (*NapA*) versus membrane (*NarG*) fractions of cell lysate showed higher activity in membrane fraction for *Alcaligenes eutrophus* H-16. | (Warnecke-Eberz and Friedrich, 1993) |

Table S2: Comparing *in vitro* nitrite reductase specific activities ($\mu\mathrm{mol}\ NO_2^- / \mathrm{min} / \mathrm{mg}$ protein) measured in previous studies.

| Organism | Reductase | Sp. act. | Ref. |
|---|---|---|---|
| *Alcaligenes xylosoxidans* NCIB 11015 | *NirK* | 244 | (Abraham et al., 1993) |
| *Alcaligenes xylosoxidans* NCIB 11015 | *NirK* | 117 | (Masuko et al., 1984) |
| *Achromobacter cyclastes* IAM 1013 | *NirK* | 149 | (Iwasaki and Matsubara, 1972) |
| *Achromobacter cyclastes* IAM 1013 | *NirK* | 280 | (Liu et al., 1986) |
| *Alcaligenes faecalis* S-6 | *NirK* | 380 | (Kakutani et al., 1981) |
| *Alcaligenes faecalis* S-6 | *NirK* | 350 | (Kukimoto et al., 1994) |
| *Rhodobacter sphaeroides* IL 106 | *NirK* | 40 | (Michalski and Nicholas, 1985) |
| *Rhodobacter sphaeroides* IL 106 | *NirK* | 26.2 | (Sawada et al., 1978) |
| *Bacillus halodenitrificans* | *NirK* | 90 | (Denariaz et al., 1991) |
| *Pseudomonas aeruginosa* PA01 | *NirS* | 3.78 | (Zumft, 1997) |
| *Pseudomonas stutzeri* ZoBell | *NirS* | 4.15 | (Zumft, 1997) |
| *Paracoccus denitrificans* ATCC 13543 | *NirS* | 4.0 | (Timkovich et al., 1982) |
| *Paracoccus pantrophus* LMD92.63 | *NirS* | 34 | (Gordon et al., 2003) |
| *Pseudomonas nautica* 617 | *NirS* | 0.048 | (Besson et al., 1995) |
| *Thiobacillus denitrificans* NCIB AB5 | *NirS* | 0.037 | (Sawhney and Nicholas, 1978) |

Table S3: The dates and locations of soil sample collection, as well as the treatments applied to each soil sample prior to isolation, and the isolates obtained from each sample.

| Soil sample | Location | Date collected | Latitude | Longitude | Treatment | Strains isolated |
|---|---|---|---|---|---|---|
| sia01 | Forest (Busey Woods) | 10/4/18 | 40.127 | -88.209 | Incubated at room temperature for 7 days | |
| sia04 | Forest (Busey Woods) | 10/4/18 | 40.127 | -88.209 | Incubated at room temperature for 7 days | |
| sia05 | Prairie (Meadowbrook) | 10/4/18 | 40.079 | -88.205 | Incubated at room temperature for 7 days | |
| sia06 | Lawn (Japan House) | 10/4/18 | 40.093 | -88.218 | Incubated at room temperature for 7 days | |
| sia07 | Agricultural (drainage ditch) | 10/4/18 | 40.085 | -88.232 | Incubated at room temperature for 7 days | PDM18 |
| sia09 | Lawn (Loomis) | 10/4/18 | 40.111 | -88.222 | Incubated at room temperature for 7 days | PDM13, PDM20 |
| sib01 | Lawn (Japan House) | 10/18/18 | 40.093 | -88.218 | Incubated at room temperature for 1 day | PDM21, RLT01 |
| sib02 | Forest (Busey Woods) | 10/18/18 | 40.127 | -88.209 | Incubated at room temperature for 1 day | |
| sib03 | Agricultural (soy field) | 10/18/18 | 40.057 | -88.232 | Incubated at room temperature for 1 day | |
| sib04 | Agricultural (corn field) | 10/18/18 | 40.057 | -88.232 | Incubated at room temperature for 1 day | PNT03, VRV01 |
| sib05 | Prairie (Meadowbrook) | 10/18/18 | 40.079 | -88.205 | Incubated at room temperature for 1 day | |
| sib06 | Lawn (Loomis) | 10/18/18 | 40.111 | -88.222 | Incubated at room temperature for 1 day | PDM12 |
| sic01 | Lawn (Japan House) | 10/18/18 | 40.093 | -88.218 | Incubated at room temperature for 1 day | |
| sic02 | Lawn (Japan House) | 10/18/18 | 40.093 | -88.218 | Incubated at room temperature for 1 day | PNT02 |
| sic03 | Forest (Busey Woods) | 10/18/18 | 40.127 | -88.209 | Incubated at room temperature for 1 day | BKH01 |
| sic04 | Forest (Busey Woods) | 10/18/18 | 40.127 | -88.209 | Incubated at room temperature for 1 day | ENS09 |
| sic05 | Agricultural (soy field) | 10/18/18 | 40.057 | -88.232 | Incubated at room temperature for 1 day | |
| sic06 | Agricultural (soy field) | 10/18/18 | 40.057 | -88.232 | Incubated at room temperature for 1 day | PDM07 |
| sic07 | Agricultural (corn field) | 10/18/18 | 40.057 | -88.232 | Incubated at room temperature for 1 day | |
| sic08 | Agricultural (corn field) | 10/18/18 | 40.057 | -88.232 | Incubated at room temperature for 1 day | |
| sic09 | Agricultural (corn field) | 10/18/18 | 40.057 | -88.232 | Incubated at room temperature for 1 day | PDM08 |
| sic10 | Agricultural (corn field) | 10/18/18 | 40.057 | -88.232 | Incubated at room temperature for 1 day | PDM01, PDM02, PDM03 |
| sic11 | Prairie (Meadowbrook) | 10/18/18 | 40.079 | -88.205 | Incubated at room temperature for 1 day | PDM10 |
| sic12 | Prairie (Meadowbrook) | 10/18/18 | 40.079 | -88.205 | Incubated at room temperature for 1 day | PDM09 |
| sic13 | Lawn (Loomis) | 10/18/18 | 40.111 | -88.222 | Incubated at room temperature for 1 day | |
| sic14 | Lawn (Loomis) | 10/18/18 | 40.111 | -88.222 | Incubated at room temperature for 1 day | |
| sic15 | Lawn (Loomis) | 10/18/18 | 40.111 | -88.222 | Incubated at room temperature for 1 day | |
| sic16 | Lawn (Loomis) | 10/18/18 | 40.111 | -88.222 | Incubated at room temperature for 1 day | PNT01 |
| sid01 | Lawn (Japan House) | 10/18/18 | 40.093 | -88.218 | Incubated at 30 C for 6 days with 10 mM NO2- | ENS08, PDM19 |
| sid02 | Lawn (Japan House) | 10/18/18 | 40.093 | -88.218 | Incubated at 30 C for 6 days with 10 mM NO2- | ENS07, PDM23 |
| sid03 | Forest (Busey Woods) | 10/18/18 | 40.127 | -88.209 | Incubated at 30 C for 6 days with 10 mM NO2- | ENS05, ENS06, STM01 |
| sid04 | Forest (Busey Woods) | 10/18/18 | 40.127 | -88.209 | Incubated at 30 C for 6 days with 10 mM NO2- | CMM01, DLF01, ENS04, PAR01 |
| sid05 | Agricultural (soy field) | 10/18/18 | 40.057 | -88.232 | Incubated at 30 C for 6 days with 10 mM NO2- | |
| sid06 | Agricultural (corn field) | 10/18/18 | 40.057 | -88.232 | Incubated at 30 C for 6 days with 10 mM NO2- | ENS03 |
| sid07 | Prairie (Meadowbrook) | 10/18/18 | 40.079 | -88.205 | Incubated at 30 C for 6 days with 10 mM NO2- | PDM17, PXM03 |
| sid08 | Prairie (Meadowbrook) | 10/18/18 | 40.079 | -88.205 | Incubated at 30 C for 6 days with 10 mM NO2- | ENS10, PDM14, PDM22 |
| sid09 | Lawn (Loomis) | 10/18/18 | 40.111 | -88.222 | Incubated at 30 C for 6 days with 10 mM NO2- | XNM01 |
| sid10 | Lawn (Loomis) | 10/18/18 | 40.111 | -88.222 | Incubated at 30 C for 6 days with 10 mM NO2- | ENS01, ENS02, ENS11, PDM16 |
| sie01 | Prairie (Meadowbrook) | 5/24/19 | 40.079 | -88.205 | No incubation | |
| sie02 | Lawn (Loomis) | 5/24/19 | 40.111 | -88.222 | No incubation | |
| sie03 | Lawn (Loomis) | 5/24/19 | 40.111 | -88.222 | No incubation | |
| sie04 | Lawn (Loomis) | 5/24/19 | 40.111 | -88.222 | No incubation | |
| sie05 | Lawn (Loomis) | 5/24/19 | 40.111 | -88.222 | No incubation | |
| sie06 | Lawn (Loomis) | 5/24/19 | 40.111 | -88.222 | No incubation | |
| sie07 | Lawn (Loomis) | 5/24/19 | 40.111 | -88.222 | No incubation | |
| sie08 | Lawn (Loomis) | 5/24/19 | 40.111 | -88.222 | No incubation | |
| sie09 | Lawn (Loomis) | 5/24/19 | 40.111 | -88.222 | No incubation | |
| sie10 | Lawn (Loomis) | 5/24/19 | 40.111 | -88.222 | No incubation | |
| sie11 | Lawn (Loomis) | 5/24/19 | 40.111 | -88.222 | No incubation | |
| sie12 | Lawn (Loomis) | 5/24/19 | 40.111 | -88.222 | No incubation | |
| sie13 | Lawn (Japan House) | 5/24/19 | 40.093 | -88.218 | No incubation | |
| sie14 | Lawn (Japan House) | 5/24/19 | 40.093 | -88.218 | No incubation | PXM02 |
| sie15 | Lawn (Japan House) | 5/24/19 | 40.093 | -88.218 | No incubation | |
| sie16 | Lawn (Japan House) | 5/24/19 | 40.093 | -88.218 | No incubation | |
| sie17 | Lawn (Japan House) | 5/24/19 | 40.093 | -88.218 | No incubation | |
| sie18 | Lawn (Japan House) | 5/24/19 | 40.093 | -88.218 | No incubation | |
| sie19 | Lawn (Japan House) | 5/24/19 | 40.093 | -88.218 | No incubation | PXM01 |
| sie20 | Lawn (Japan House) | 5/24/19 | 40.093 | -88.218 | No incubation | |
| sie21 | Lawn (Japan House) | 5/24/19 | 40.093 | -88.218 | No incubation | PDM04 |
| sie22 | Lawn (Japan House) | 5/24/19 | 40.093 | -88.218 | No incubation | |
| sie23 | Lawn (Japan House) | 5/24/19 | 40.093 | -88.218 | No incubation | |
| sie24 | Lawn (Japan House) | 5/24/19 | 40.093 | -88.218 | No incubation | |
| sif01 | Prairie (Meadowbrook) | 5/24/19 | 40.079 | -88.205 | Incubated at 30 C for 14 days with 1 mM NO2- | |
| sif02 | Lawn (Loomis) | 5/24/19 | 40.111 | -88.222 | Incubated at 30 C for 14 days with 1 mM NO2- | |
| sif03 | Lawn (Loomis) | 5/24/19 | 40.111 | -88.222 | Incubated at 30 C for 14 days with 1 mM NO2- | |
| sif04 | Lawn (Loomis) | 5/24/19 | 40.111 | -88.222 | Incubated at 30 C for 14 days with 1 mM NO2- | |
| sif05 | Lawn (Loomis) | 5/24/19 | 40.111 | -88.222 | Incubated at 30 C for 14 days with 1 mM NO2- | ACM01, PDM06, PXM04 |
| sif06 | Lawn (Loomis) | 5/24/19 | 40.111 | -88.222 | Incubated at 30 C for 14 days with 1 mM NO2- | ACM03, ARZ01, PDM05 |
| sif07 | Lawn (Loomis) | 5/24/19 | 40.111 | -88.222 | Incubated at 30 C for 14 days with 1 mM NO2- | AGB01, PDM15 |
| sif08 | Lawn (Loomis) | 5/24/19 | 40.111 | -88.222 | Incubated at 30 C for 14 days with 1 mM NO2- | |
| sif09 | Lawn (Loomis) | 5/24/19 | 40.111 | -88.222 | Incubated at 30 C for 14 days with 1 mM NO2- | |
| sif10 | Lawn (Loomis) | 5/24/19 | 40.111 | -88.222 | Incubated at 30 C for 14 days with 1 mM NO2- | |
| sif11 | Lawn (Loomis) | 5/24/19 | 40.111 | -88.222 | Incubated at 30 C for 14 days with 1 mM NO2- | |
| sif12 | Lawn (Loomis) | 5/24/19 | 40.111 | -88.222 | Incubated at 30 C for 14 days with 1 mM NO2- | |
| sif13 | Lawn (Japan House) | 5/24/19 | 40.093 | -88.218 | Incubated at 30 C for 14 days with 1 mM NO2- | |
| sif14 | Lawn (Japan House) | 5/24/19 | 40.093 | -88.218 | Incubated at 30 C for 14 days with 1 mM NO2- | |
| sif15 | Lawn (Japan House) | 5/24/19 | 40.093 | -88.218 | Incubated at 30 C for 14 days with 1 mM NO2- | |
| sif16 | Lawn (Japan House) | 5/24/19 | 40.093 | -88.218 | Incubated at 30 C for 14 days with 1 mM NO2- | RHZ01 |
| sif17 | Lawn (Japan House) | 5/24/19 | 40.093 | -88.218 | Incubated at 30 C for 14 days with 1 mM NO2- | ACM02, RHZ02 |
| sif18 | Lawn (Japan House) | 5/24/19 | 40.093 | -88.218 | Incubated at 30 C for 14 days with 1 mM NO2- | CMM02, ENT01 |
| sif19 | Lawn (Japan House) | 5/24/19 | 40.093 | -88.218 | Incubated at 30 C for 14 days with 1 mM NO2- | |
| sif20 | Lawn (Japan House) | 5/24/19 | 40.093 | -88.218 | Incubated at 30 C for 14 days with 1 mM NO2- | |
| sif21 | Lawn (Japan House) | 5/24/19 | 40.093 | -88.218 | Incubated at 30 C for 14 days with 1 mM NO2- | |
| sif22 | Lawn (Japan House) | 5/24/19 | 40.093 | -88.218 | Incubated at 30 C for 14 days with 1 mM NO2- | ACV02, PDM11 |
| sif23 | Lawn (Japan House) | 5/24/19 | 40.093 | -88.218 | Incubated at 30 C for 14 days with 1 mM NO2- | ACV01 |
| sif24 | Lawn (Japan House) | 5/24/19 | 40.093 | -88.218 | Incubated at 30 C for 14 days with 1 mM NO2- | ACM04 |

Table S4: Components and final concentrations of the succinate defined medium (SDM).

| Name | Concentration |
|---|---|
| $Na_2HPO_4 \cdot 7\ H_2O$ | 32.01 mM |
| Sodium succinate dibasic $\cdot$ 7 $H_2O$ | 25.00 mM |
| $KH_2PO_4$ | 8.01 mM |
| $(NH_4)_2SO_4$ | 7.57 mM |
| $MgSO_4$ | 2.40 mM |
| Nitrilotriacetic acid | 1.05 mM |
| $CaCl_2 \cdot 2\ H_2O$ | 0.45 mM |
| $ZnSO_4 \cdot 7\ H_2O$ | 38.25 µM |
| $FeSO_4 \cdot 7\ H_2O$ | 25.11 µM |
| $MnSO_4 \cdot H_2O$ | 9.11 µM |
| $Na_2 \cdot EDTA \cdot 2\ H_2O$ | 8.55 µM |
| $CuSO_4 \cdot 5\ H_2O$ | 1.57 µM |
| $CoCl_2 \cdot 6H_2O$ | 0.85 µM |
| Nicotinic acid | 0.81 µM |
| Pyridoxine hydrochloride | 0.73 µM |
| $Na_2B_4O_7 \cdot 10\ H_2O$ | 0.46 µM |
| Thiamine HCL | 0.30 µM |
| 4-aminobenzoic acid | 0.29 µM |
| Riboflavin | 0.27 µM |
| $(NH_4)_6Mo_7O_{24} \cdot 4\ H_2O$ | 0.15 µM |
| Calcium D(+) pantothenate | 0.10 µM |
| Folic acid | 90.62 nM |
| D-(+) biotin | 81.86 nM |
| Cyanocobalamin | 73.78 nM |
| Lipoic acid | 48.47 nM |

Table S5: Time to saturation for the $62$-strain library when grown aerobically on SDM at $30\,°C$. Cultures were inoculated from saturated pre-cultures grown aerobically in 1/5X TSB.

| Strain | Saturation time (h) |
|---|---|
| ACM01 | 14.5 |
| ACM02 | 14.5 |
| ACM03 | 14.8 |
| ACM04 | 14.8 |
| ACV01 | 21.5 |
| ACV02 | 14.0 |
| AGB01 | 23.0 |
| ARZ01 | 26.4 |
| BKH01 | 18.0 |
| CMM01 | 17.8 |
| CMM02 | 15.8 |
| DLF01 | 12.8 |
| ENS01 | 22.8 |
| ENS02 | 18.7 |
| ENS03 | 18.8 |
| ENS04 | 23.3 |
| ENS05 | 20.2 |
| ENS06 | 17.5 |
| ENS07 | 18.7 |
| ENS08 | 18.7 |
| ENS09 | 19.8 |
| ENS10 | 20.8 |
| ENS11 | 19.5 |
| ENT01 | 72.3 |
| PAR01 | 15.8 |
| PAR19367 | 11.5 |
| PDM01 | 14.8 |
| PDM02 | 15.3 |
| PDM03 | 16.0 |
| PDM04 | 15.5 |
| PDM05 | 14.8 |
| PDM06 | 14.8 |
| PDM07 | 14.3 |
| PDM08 | 14.5 |
| PDM09 | 16.3 |
| PDM10 | 18.8 |
| PDM11 | 12.5 |
| PDM12 | 13.0 |
| PDM13 | 13.0 |
| PDM14 | 19.0 |
| PDM15 | 36.5 |
| PDM16 | 14.5 |
| PDM17 | 13.5 |
| PDM18 | 11.5 |
| PDM19 | 13.5 |
| PDM20 | 13.2 |
| PDM21 | 13.0 |
| PDM22 | 13.3 |
| PDM23 | 13.5 |
| PNT01 | 21.5 |
| PNT02 | 17.0 |
| PNT03 | 14.8 |
| PXM01 | 51.8 |
| PXM02 | 51.8 |
| PXM03 | 44.3 |
| PXM04 | 28.3 |
| RHZ01 | 21.0 |
| RHZ02 | 23.0 |
| RLT01 | 62.8 |
| STM01 | 25.0 |
| VRV01 | 19.8 |
| XNM01 | 18.7 |

Table S6: Gene function labels identifying main denitrifying genes (i.e., terminal reductases, sensors/regulators/transporters) in RAST annotations.

| RAST label | Gene name |
|---|---|
| Respiratory nitrate reductase alpha chain (EC 1.7.99.4) | narG |
| Periplasmic nitrate reductase (EC 1.7.99.4) | napA |
| Copper-containing nitrite reductase (EC 1.7.2.1) | nirK |
| Nitrite reductase (EC 1.7.2.1) | nirS |
| Nitric-oxide reductase subunit B (EC 1.7.99.7) | cnor |
| Nitric-oxide reductase (EC 1.7.99.7), quinol-dependent | qnor |
| Nitrous-oxide reductase (EC 1.7.99.6) | nosZ |
| Nitrate/nitrite sensor protein NarX | narXL |
| Nitrate/nitrite sensor protein | narXL |
| Nitrate/nitrite sensor protein (EC 2.7.3.-) | narXL |
| Nitrate/nitrite response regulator protein NarL | narXL |
| Nitrate/nitrite response regulator protein | narXL |
| Nitrate/nitrite sensor protein NarQ | narXL |
| Nitrate/nitrite response regulator protein NarP | narXL |
| Nitric oxide -responding transcriptional regulator Dnr (Crp/Fnr family) | dnr |
| Nitric oxide -responding transcriptional regulator NnrR (Crp/Fnr family) | nnrR |
| NnrS protein involved in response to NO | nnrS |
| NnrU family protein, required for expression of nitric oxide and nitrite reductases (Nir and Nor) | nnrU |
| Nitrite-sensitive transcriptional repressor NsrR | nsrR |
| Anaerobic nitric oxide reductase transcription regulator NorR | norR |
| Nitrate/nitrite transporter NarK/U 1 | narK1 |
| Nitrate/nitrite transporter NarK/U | narK2 |
| Nitrate/nitrite transporter NarU | narK2 |
| Nitrate/nitrite transporter NarK/U 1 / Nitrate/nitrite transporter NarK/U | narK1K2 |

Table S7: Gene function labels identifying auxiliary denitrifying genes (e.g., chaperones, structural genes, and biosynthesis genes that frequently occur in clusters with the terminal reductases) in RAST annotations.

| RAST label | Gene name |
|---|---|
| Respiratory nitrate reductase beta chain (EC 1.7.99.4) | narH |
| Respiratory nitrate reductase gamma chain (EC 1.7.99.4) | narI |
| Respiratory nitrate reductase delta chain (EC 1.7.99.4) | narJ |
| Nitrate reductase cytochrome c550-type subunit | napB |
| Cytochrome c-type protein NapC | napC |
| Periplasmic nitrate reductase component NapD | napD |
| Periplasmic nitrate reductase component NapE | napE |
| Ferredoxin-type protein NapF (periplasmic nitrate reductase) | napF |
| Cytochrome c55X precursor NirC | nirC |
| Heme d1 biosynthesis protein NirF | nirF |
| Heme d1 biosynthesis protein NirG | nirG |
| Heme d1 biosynthesis protein NirH | nirH |
| Heme d1 biosynthesis protein NirJ | nirJ |
| Heme d1 biosynthesis protein NirL | nirL |
| Nitrite reductase associated c-type cytochorome NirN | nirN |
| Nitrite reductase accessory protein NirV | nirV |
| Nitric-oxide reductase subunit C (EC 1.7.99.7) | norC |
| Nitric oxide reductase activation protein NorD | norD |
| Nitric oxide reductase activation protein NorE | norE |
| Nitric oxide reductase activation protein NorQ | norQ |
| Nitrous oxide reductase maturation protein NosD | nosD |
| Nitrous oxide reductase maturation protein NosF (ATPase) | nosF |
| Nitrous oxide reductase maturation protein, outer-membrane lipoprotein NosL | nosL |
| Nitrous oxide reductase maturation protein NosR | nosR |
| Nitrous oxide reductase maturation transmembrane protein NosY | nosY |

Table S8: Estimated genome sizes and 16S rRNA copy numbers for each strain in the $62$-strain library.

| Strain | Genome size (mb) | 16S rRNA copy number |
|---|---|---|
| ACM01 | 6.63 | 4.04 |
| ACM02 | 6.49 | 3.39 |
| ACM03 | 6.72 | 3.98 |
| ACM04 | 6.62 | 3.93 |
| ACV01 | 5.85 | 2.97 |
| ACV02 | 5.20 | 4.17 |
| AGB01 | 5.62 | 5.22 |
| ARZ01 | 5.21 | 3.05 |
| BKH01 | 7.62 | 5.48 |
| CMM01 | 4.72 | 5.85 |
| CMM02 | 3.89 | 6.87 |
| DLF01 | 6.43 | 4.62 |
| ENS01 | 7.37 | 5.35 |
| ENS02 | 7.50 | 4.66 |
| ENS03 | 7.21 | 4.86 |
| ENS04 | 7.74 | 4.93 |
| ENS05 | 7.63 | 4.80 |
| ENS06 | 7.92 | 5.26 |
| ENS07 | 7.95 | 4.86 |
| ENS08 | 7.31 | 4.53 |
| ENS09 | 7.02 | 4.85 |
| ENS10 | 7.65 | 4.28 |
| ENS11 | 7.34 | 5.08 |
| ENT01 | 4.62 | 6.98 |
| PAR01 | 5.11 | 4.95 |
| PAR19367 | 5.15 | 3.36 |
| PDM01 | 6.36 | 5.96 |
| PDM02 | 6.40 | 6.25 |
| PDM03 | 6.53 | 5.16 |
| PDM04 | 6.46 | 4.48 |
| PDM05 | 6.03 | 5.98 |
| PDM06 | 7.01 | 4.73 |
| PDM07 | 6.50 | 5.39 |
| PDM08 | 6.56 | 5.85 |
| PDM09 | 6.34 | 5.71 |
| PDM10 | 6.52 | 6.78 |
| PDM11 | 4.67 | 3.94 |
| PDM12 | 5.18 | 4.26 |
| PDM13 | 6.66 | 4.00 |
| PDM14 | 4.65 | 3.12 |
| PDM15 | 4.84 | 2.92 |
| PDM16 | 5.09 | 4.18 |
| PDM17 | 6.64 | 4.03 |
| PDM18 | 6.43 | 5.16 |
| PDM19 | 6.81 | 4.72 |
| PDM20 | 6.69 | 4.65 |
| PDM21 | 6.75 | 5.03 |
| PDM22 | 6.71 | 4.94 |
| PDM23 | 6.70 | 4.19 |
| PNT01 | 4.58 | 6.29 |
| PNT02 | 5.20 | 6.74 |
| PNT03 | 4.97 | 6.80 |
| PXM01 | 4.18 | 2.09 |
| PXM02 | 4.34 | 1.94 |
| PXM03 | 4.04 | 2.37 |
| PXM04 | 4.29 | 2.11 |
| RHZ01 | 6.43 | 4.28 |
| RHZ02 | 6.52 | 3.98 |
| RLT01 | 5.36 | 7.70 |
| STM01 | 4.14 | 3.63 |
| VRV01 | 7.02 | 2.09 |
| XNM01 | 4.51 | 2.02 |

Table S9: 12-strain subset of the 62-library used for community assembly experiments.

| Strain label | Class | Genus | Phenotype |
|---|---|---|---|
| ACM01 | Betaproteobacteria | *Achromobacter* | Nar/Nir |
| ENS09 | Alphaproteobacteria | *Ensifer* | Nar/Nir |
| PAR19367 | Alphaproteobacteria | *Paracoccus* | Nar/Nir |
| PDM21 | Gammaproteobacteria | *Pseudomonas* | Nar/Nir |
| ACV02 | Betaproteobacteria | *Acidovorax* | Nar |
| PAR01 | Alphaproteobacteria | *Paracoccus* | Nar |
| PDM12 | Gammaproteobacteria | *Pseudomonas* | Nar |
| PNT03 | Gammaproteobacteria | *Pantoea* | Nar |
| AGB01 | Alphaproteobacteria | *Agrobacterium* | Nir |
| PDM13 | Gammaproteobacteria | *Pseudomonas* | Nir |
| PDM14 | Gammaproteobacteria | *Pseudomonas* | Nir |
| PXM03 | Gammaproteobacteria | *Pseudoxanthomonas* | Nir |
